# Identification of RNA 3’ ends and termination sites in *Haloferax volcanii*

**DOI:** 10.1101/748780

**Authors:** Sarah J. Berkemer, Lisa-Katharina Maier, Fabian Amman, Stephan H. Bernhart, Julia Wörtz, Pascal Märkle, Friedhelm Pfeiffer, Peter F. Stadler, Anita Marchfelder

**Affiliations:** Bioinformatics Group, Department of Computer Science - and Interdisciplinary Center for Bioinformatics, University of Leipzig, Härtelstraße 16-18, D-04107 Leipzig, Germany; Max Planck Institute for Mathematics in the Sciences, Inselstraße 22, D-04103 Leipzig, Germany; Biology II, Ulm University, 89069 Ulm, Germany; Computational Biology Group, Max Planck Institute of Biochemistry, 82152 Martinsried, Germany; Transcriptome Bioinformatics, Interdisciplinary Center for Bioinformatics, Leipzig University, Germany; German Centre for Integrative Biodiversity Research (iDiv) Halle-Jena-Leipzig; Competence Center for Scalable Data Services and Solutions, and Leipzig, Research Center for Civilization Diseases, University Leipzig, Germany; Facultad de Ciencias, Universidad Nacional de Colombia, Bogotá, Colombia; Institute for Theoretical Chemistry, University of Vienna, Währingerstraße 17, A-1090 Vienna, Austria; Division of Cell and Developmental Biology, Medical University Vienna, Schwarzspanierstraße 17, 1090 Vienna, Austria; Center for RNA in Technology and Health, Univ. Copenhagen, Grønnegårdsvej 3, Frederiksberg C, Denmark; Santa Fe Institute, 1399 Hyde Park Rd., Santa Fe, NM 87501

## Abstract

Archaeal genomes are densely packed; thus, correct transcription termination is an important factor for orchestrated gene expression. A systematic analysis of RNA 3’ termini, to identify transcription termination sites (TTS) using RNAseq data has hitherto only been performed in two archaea. In this study, only part of the genome had been investigated. Here, we developed a novel algorithm that allows an unbiased, genome-wide identification of RNA 3’ termini independent of annotation. In an RNA fraction enriched for primary transcripts by terminator exonuclease (TEX) treatment we identified 1,543 RNA 3’ termini. A strong sequence signature consistent with known termination events at intergenic loci indicates a clear enrichment for native TTS among them. Using these data we determined distinct putative termination motifs for intergenic (a T stretch) and coding regions (AGATC). In vivo reporter gene tests of selected TTS confirmed termination at these sites, which exemplify the different motifs. For several genes, more than one termination site was detected, resulting in transcripts with different lengths of the 3’ untranslated region.

## Introduction

Archaeal RNA synthesis is generally considered to be more closely related to transcription in eukaryotes than to bacterial transcription. The archaeal RNA polymerase is similar to the eukaryotic RNA polymerase II, and the basal promoter elements in Archaea are similar to their eukaryotic pendants (TATA box and BRE); general transcription factors TBP (TATA binding protein), TFB (transcription factor B) and TFE (transcription factor E) resemble the eukaryotic proteins TBP, TFIIB and TFEII, respectively (for a review see: Fouqueau et al. (1)). Transcriptional regulators, however, seem to be more similar to those in bacteria (2). Thus, the archaeal transcription machinery consists of a mixture of bacterial-like and eukaryotic-like components. Whereas some data have been reported on transcription initiation and elongation in archaea, very little is known about transcription termination. Controlled transcription termination is important to avoid aberrant RNA molecules and to help with RNA polymerase recycling. Generally, the genes in archaeal chromosomes are densely packed, so that proper termination is also important to prevent transcription from continuing into downstream genes. The process of transcription termination is not trivial because the very stable transcription elongation complex must be destabilised and dissociated during termination. In bacteria, two major classes of termination signals have been described: intrinsic termination and factor-dependent termination (3,4). Intrinsic termination occurs either at a stretch of Ts or at hairpin structures that fold in the newly synthesised RNA; both trigger dissociation of the elongation complex. Factor-dependent termination occurs upon interaction with a specific protein such as the bacterial termination factor Rho (5). Protein factor-assisted termination is especially important in regions where strong selective pressure on the DNA sequence does not allow encoding of intrinsic termination signals. This may be the case when termination must occur in the coding region of a downstream gene. In eukaryotes, several RNA polymerases synthesise the different RNA classes, and these polymerases have different modes of termination. RNA polymerase I requires protein factors for termination (6), whereas RNA polymerase III terminates efficiently and precisely at a stretch of Ts (7). Termination of RNA polymerase II is quite complex, involving modification of the RNAP as well as interactions with additional protein factors, and seems to be coupled to co-transcriptional RNA processing (8). Whereas archaeal transcription initiation resembles eukaryotic RNA polymerase II initiation (9), transcription elongation and termination seem to be more similar to the eukaryotic RNA polymerase III pathway, which is independent of RNA secondary structures and protein co-factors (10).

Compared to the determination of transcription start sites, the identification of termination sites is more complex. Termination is often leaky and encompasses several consecutive sites, and degradation by exonucleases renders the 3’ ends heterogeneous and less clear. Also, original transcription termination sites are difficult to distinguish from RNA 3’ termini that are due to RNA processing. Data reported on archaeal termination so far show that intrinsic termination occurs at a run of Ts (10–14) and is potentially also influenced by secondary structure elements (10,14). Factor-dependent termination was predicted based on the results of an *in vivo* reporter assay in the archaeon *Thermococcus kodakarensis* (15). A recent study confirmed this hypothesis, reporting the discovery of the first archaeal termination factor (16).

Recently, RNA 3’ ends for two archaea (*Sulfolobus acidocaldarius* and *Methanosarcina mazei*) were investigated systematically using RNAseq data (17). All identified RNA 3’ ends were considered to reflect transcription termination sites (TTS) and TTS were found for 707 and 641 transcriptional units in *S. acidocaldarius* and *M. mazei*, respectively. For more than a third of the genes analysed, multiple consecutive terminators were identified, resulting in 3’ untranslated regions (UTRs) with different lengths (17). In some cases, the terminator of an early gene in an operon was shown to be located in the downstream gene, allowing gene-specific regulation within an operon. The study also revealed lineage-specific features of termination for both archaea, confirming the requirement that more data from different archaeal organisms are required to learn more about transcription termination in this domain. However, the applied algorithm analysed only regions directly downstream of annotated genes, and thus only a part of the genome was considered. Total cellular RNA was used for the analyses and no attempt was made to distinguish between RNA 3’ ends originating from transcription termination and RNA processing.

Here, we describe the identification of RNA 3’ ends of the halophilic model archaeon *Haloferax volcanii* and the subsequent determination of termination motifs*. H. volcanii* has been used for a plethora of biological studies (18,19), including the determination of nucleosome coverage (20) and a genome-wide identification of TSS (21). *H. volcanii* requires high salt concentrations for optimal growth, and due to the high intracellular salt concentrations, RNA-protein interactions -including modes of transcription termination-may differ from those in mesophilic archaea.

We used a newly developed algorithm to identify RNA 3’ ends from RNAseq data genome-wide in an unbiased manner, independent of annotation and on the basis of reads enriched for primary transcripts. Applying this algorithm to RNAseq data from a terminator exonuclease (TEX) treated library, we found 1,543 RNA 3’ ends for the *Haloferax* genome. Subsequent analysis of the respective sites revealed a strong sequence signature consistent with the current archaeal termination model, indicating a strong enrichment of native transcription termination sites (TTS) among the newly identified RNA 3’ ends. Therefore, identified RNA 3’ ends were considered putative TTS for successive analysis with respect to primary and/or secondary structure motifs as termination signals and 3’ UTR characterisation. Selected termination motifs were confirmed using an *in vivo* reporter gene system.

## Results

### Identification of RNA 3’ ends downstream of annotated regions

To identify RNA 3’ ends downstream of annotated regions in *H. volcanii* we applied a self-implemented version of the recently published method from Dar et al. (17), which will be referred to as the Dar-Sorek-Method (DSM). RNA was isolated from *H. volcanii* cells (from three biological replicates) and cDNA libraries were generated to allow determination of RNA 3’ ends. Libraries were made such that the original RNA 3’ end is tagged and can be identified in the resulting sequence. Libraries were subjected to paired-end next-generation sequencing (NGS), resulting in 49 million reads for each library on average. 89-98% of the reads obtained mapped to the *Haloferax* genome (Supplementary Table 10). Since total RNA was used for library preparation the identified RNA 3’ ends result from either transcription termination or processing. Mapped reads were analysed with DSM, which identifies RNA 3’ ends in a defined region downstream of annotated genes at the position with the highest coverage of mapped read ends. The length of the downstream region is determined by the average insert length of corresponding paired-end reads (Figure 1).

**Figure 1.**
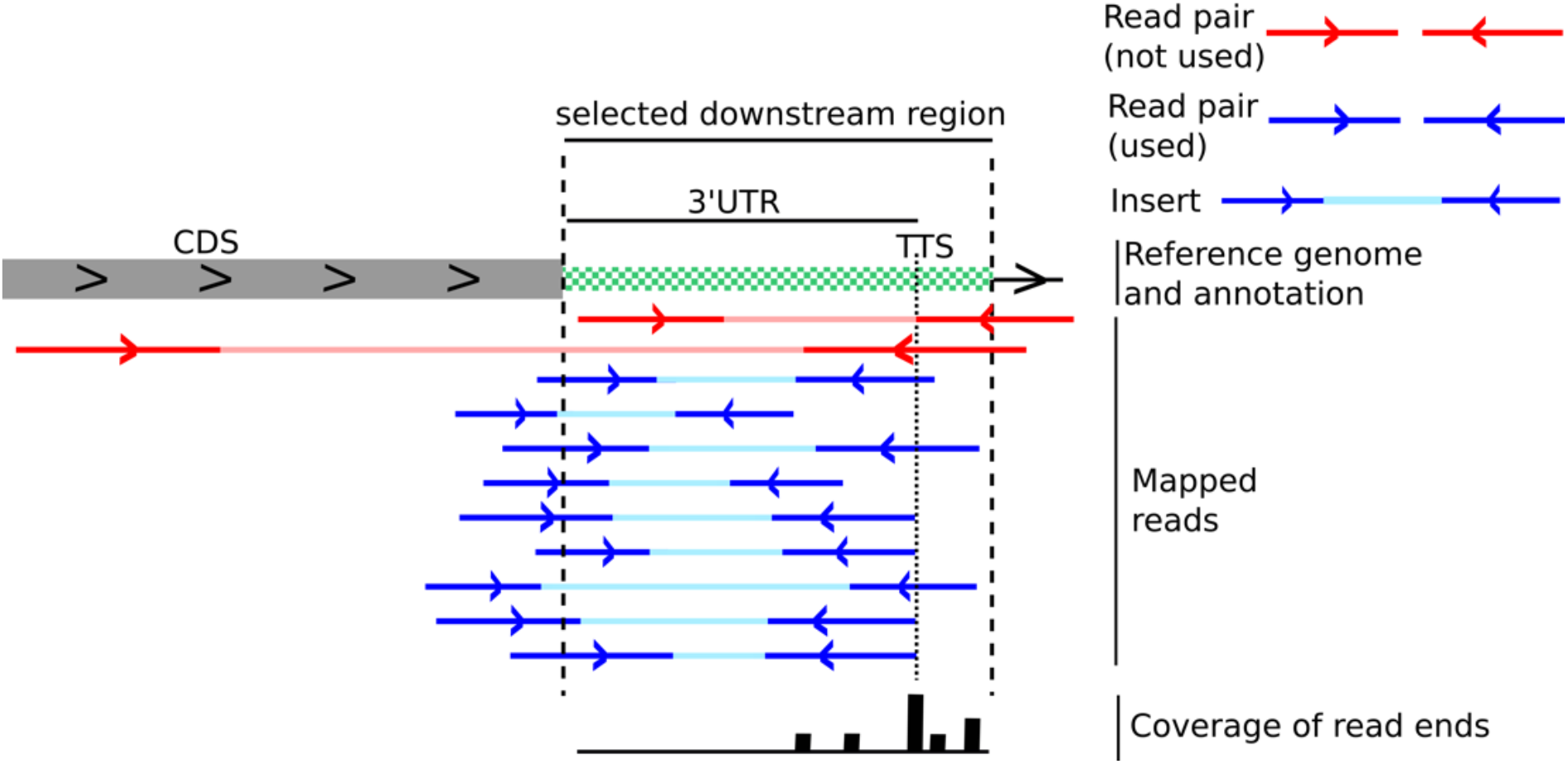
Principle of the Dar-Sorek-Method. As a first step in the DSM method, read pairs with an insert overlapping an annotated region are selected (red and blue lines)(17). Inserts that do not overlap or have a length of more than 500 nucleotides are discarded (red read pairs in the Figure). From the selected read pairs (blue), the average length over all inserts is calculated as described in Dar et al.(17). This value is used to determine the length of the selected downstream region (green white region in the Figure). The coverage of all read ends in this region is retrieved (bottom line), and the position with the highest coverage is identified as the TTS.

In our dataset, the median length of the analysed region was 126 bp. The resulting 3’ UTRs were mostly shorter than 100 nucleotides, with a median length of 58 nucleotides (Supplementary Figure 1). The length restriction given in the DSM approach might be too strict for genes with long 3’ UTRs. In addition, the method cannot determine RNA 3’ ends independent of an annotation (see also the paragraph below "Comparison between DSM and IE-PC algorithms"). Using DSM, we identified 3,155 RNA 3’ ends for the complete *Haloferax* genome of which 85% were in intergenic and the remainder in coding regions (Supplementary Table 3 and 6). A typical RNA 3’ end is shown in Supplementary Figure 2 for the *pilA2* gene (HVO_2062).

### Development of a novel algorithm, "Internal Enrichment-Peak Calling", as a tool to identify RNA 3’ ends genome-wide

DSM analysis only includes sequences downstream of annotated genes and, thus, only a fraction of the genome (17). To overcome this restriction, we developed a novel approach to interpret the RNAseq data obtained which we termed Internal Enrichment-Peak Calling (IE-PC). To this end, we utilised the nature of the fragmentation process in the course of library preparation, which leads to an enrichment of natural 3’ fragment ends over 5’ ends of fragments that are generated via random fragmentation during library preparation. Since we used the latter as the normalisation background for the former, we called this method internal enrichment (IE) (Figure 2). This approach requires mate-pair sequencing data, from which a set of read-pairs is selected that contains the same original first-strand cDNA 5’ ends (which reflects the RNA 3’ end) (Figure 2). The corresponding 3’ ends of the mate-pair reads are then evaluated. Sequencing the cDNAs generated from these first-strand cDNA fragments in paired-end mode preserved information about fragment ends. After mapping the read-pairs to the reference genome, the hallmark of an original RNA 3’ end was its high coverage of read ends, with its associated mate ends originating from a multitude of genomic sites. It is highly unlikely that independent clones end at an identical position (Figure 2). Multiple fragments with a heterogeneous 5’ end but a common 3’ end thus are indicative of true natural RNA 3’ ends.

**Figure 2.**
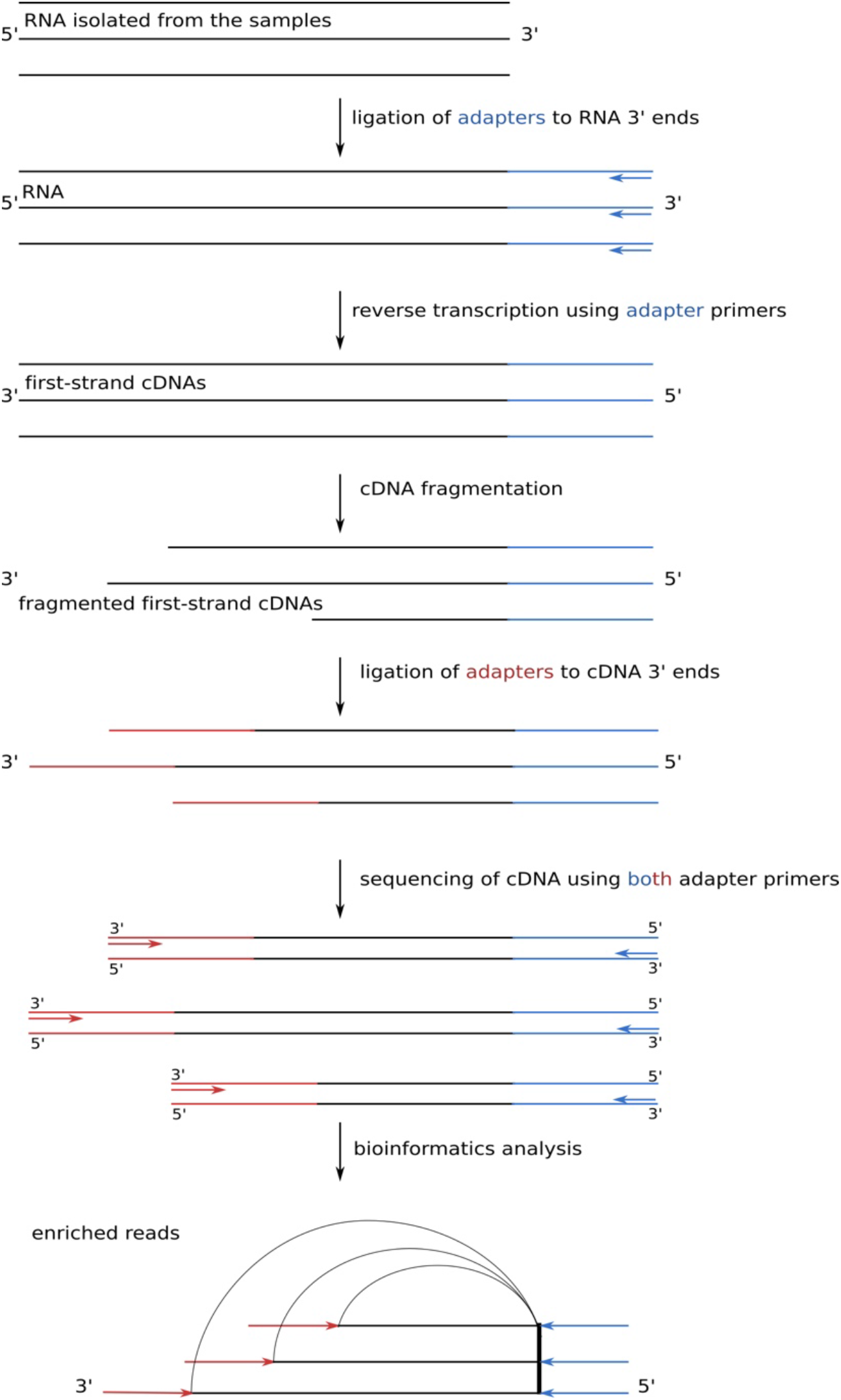
Principle of internal enrichment. After RNA isolation, adapter primers were ligated to the RNA 3’ end and the RNA was reverse transcribed. First-strand single-stranded cDNA was fragmented prior to the addition of the adapter primer at the cDNA 3’ end (for details, see Materials and Methods and Supplementary Methods). The break points were considered to be random, leading to an enrichment of original RNA 3’ ends over 5’ fragmentation ends. Sequencing the cDNAs generated from these first-strand cDNA fragments in paired-end mode preserved information about fragment ends, even if they were longer than the read length. After mapping the read-pairs to the reference genome, the hallmark of an original 3’ end was its high coverage of read ends, with its associated mate ends originating from a multitude of genomic sites.

To gain further specificity, putative 3’ ends were subsequently censored if the site in question was associated with a sudden decrease in read coverage, indicative of a true RNA 3’ end. This was assessed by a peak calling approach (termed PC), considering the absolute and relative number of fragments ending at the respective position (Figure 3). The two methods (IE and PC) were sequentially run on the data, and only sites that were found by both independent approaches, IE as well as PC, were considered to be bona-fide RNA 3’ ends. We allowed a maximal distance of 10 nucleotides when computing the intersection of IE and PC. The advantage of this algorithm is that it is independent of genome annotation and thus analyses the complete genome sequence rather than a restricted annotation-dependent region allowing the identification of all RNA 3’ ends of a genome.

**Figure 3.**
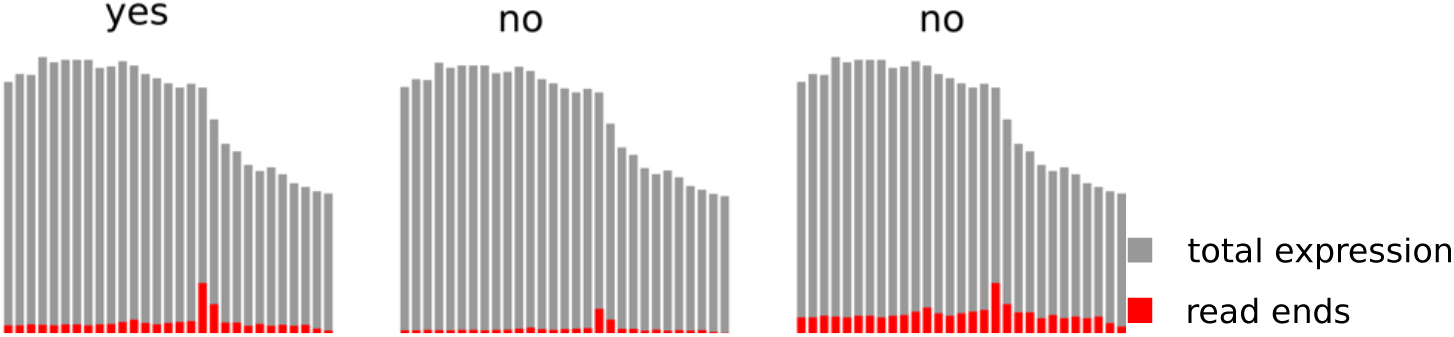
Principle of the applied peak calling procedure. In a sliding window approach, positions where the number of read stops (red) exceeded the mean number of read stops over the whole window by a z-score above the threshold 2 were treated as potential endings (left), while potential peaks of the same height but below a z-score of 2 were discarded (right). In addition, if the number of read stops was below a 10% threshold relative to the total coverage (grey), the peak was discarded (centre).

**Figure 4.**
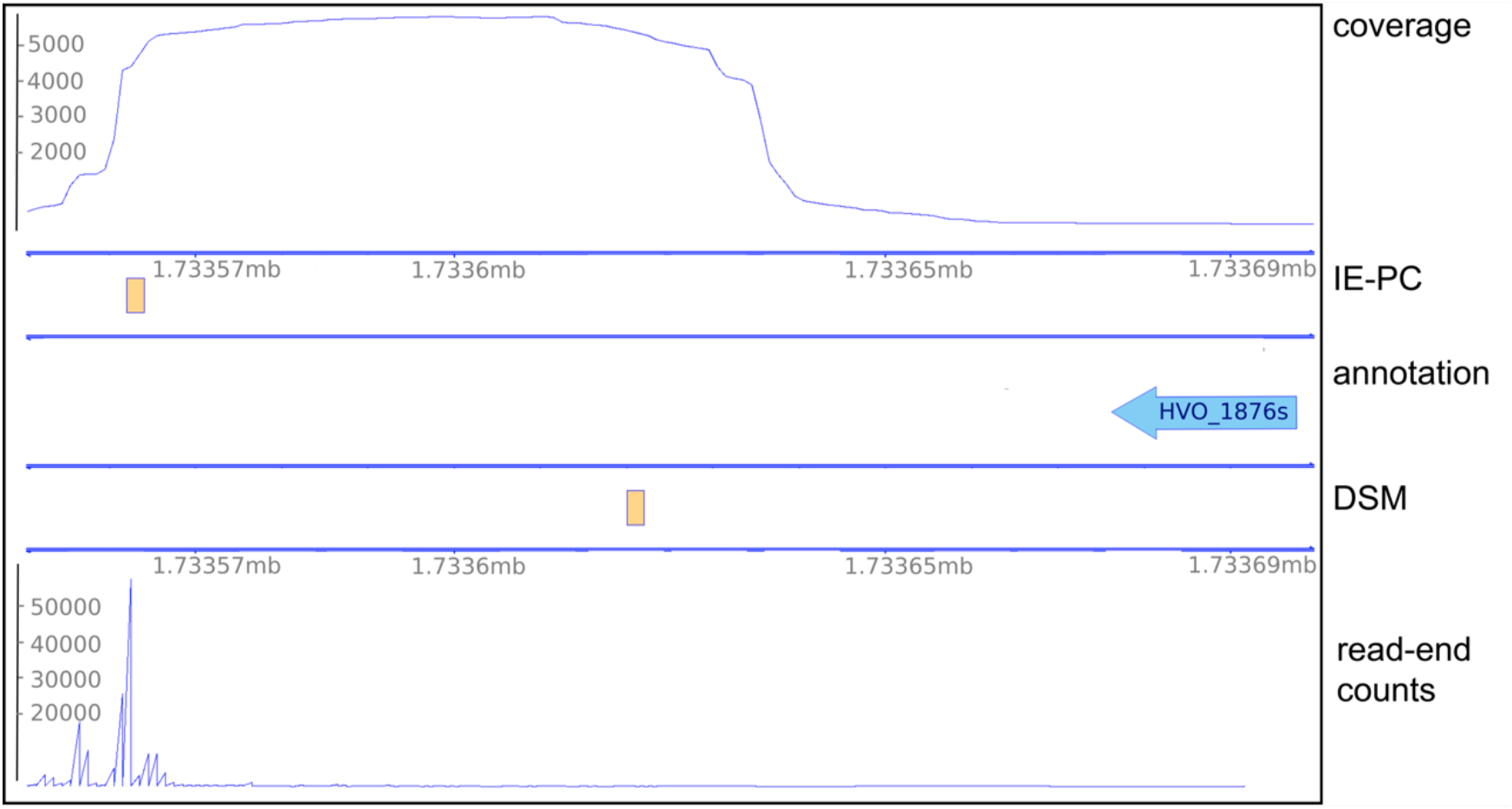
TTS comparison for DSM and IE-PC analyses for the transcript HVO_1876s. The data in the lower panel (DSM) show the DSM results with TTS and corresponding read end counts from the DSM data set. The upper panel (IE-PC) shows the TTS locations determined by IE-PC and the total coverages, corresponding to the -TEX data set. Since sequencing starts at the 3’ end, coverage starts at the 3’ end and runs continuously for 75 bp due to the read length. The annotation and genome coordinates are shown in the middle (coordinates given in Mb). TTS are shown as orange rectangles.

### Comparison between DSM and IE-PC algorithms

Since our newly established method IE-PC works independent of any genome annotation, it is able to determine RNA 3’ ends covering the complete genome. Comparison of signals obtained with both methods (DSM and IE-PC) can only be done with the regions that are included in the DSM analysis. The original DSM analysis was based on regions with a median length of 126 nucleotides downstream of an annotated 3’ end, with the RNA 3’ end located at the position with the highest coverage.

We compared individual RNA 3’ ends found with DSM and/or IE-PC in downstream regions defined by the DSM analysis in more detail. Examples are shown in Figures 5 and 6, Table 1 lists the number of RNA 3’ ends that were identified by both methods (allowing for a position discrepancy of up to 10 nucleotides) as well as RNA 3’ ends that appeared in only one of the data sets. Altogether, we found 1,664 RNA 3’ ends being present in both data sets (Table 1).

**Table 1.** Comparison of RNA 3’ ends found with DSM and IE-PC in downstream regions covered by DSM. The column "IE-PC only" lists all RNA 3’ ends identified by IE-PC but not with those identified by DSM method. RNA 3’ ends identified with IE-PC that were also found with DSM are listed in the column "overlapping". Sites that were detected in the defined region only by DSM and that did not overlap with IE-PC sites are listed in the column “DSM only”. The IE-PC data shown here are the ones calculated on the basis of the -TEX data set, since the DSM data are also based on the -TEX data.

| chromosome | IE-PC only | overlapping | DSM only |
| --- | --- | --- | --- |
| main | 3,184 | 1,296 | 1,128 |
| pHV1 | 231 | 53 | 39 |
| pHV3 | 394 | 97 | 125 |
| pHV4 | 811 | 218 | 199 |
| **total** | **4,620** | **1,664** | **1,491** |

**Figure 5.**
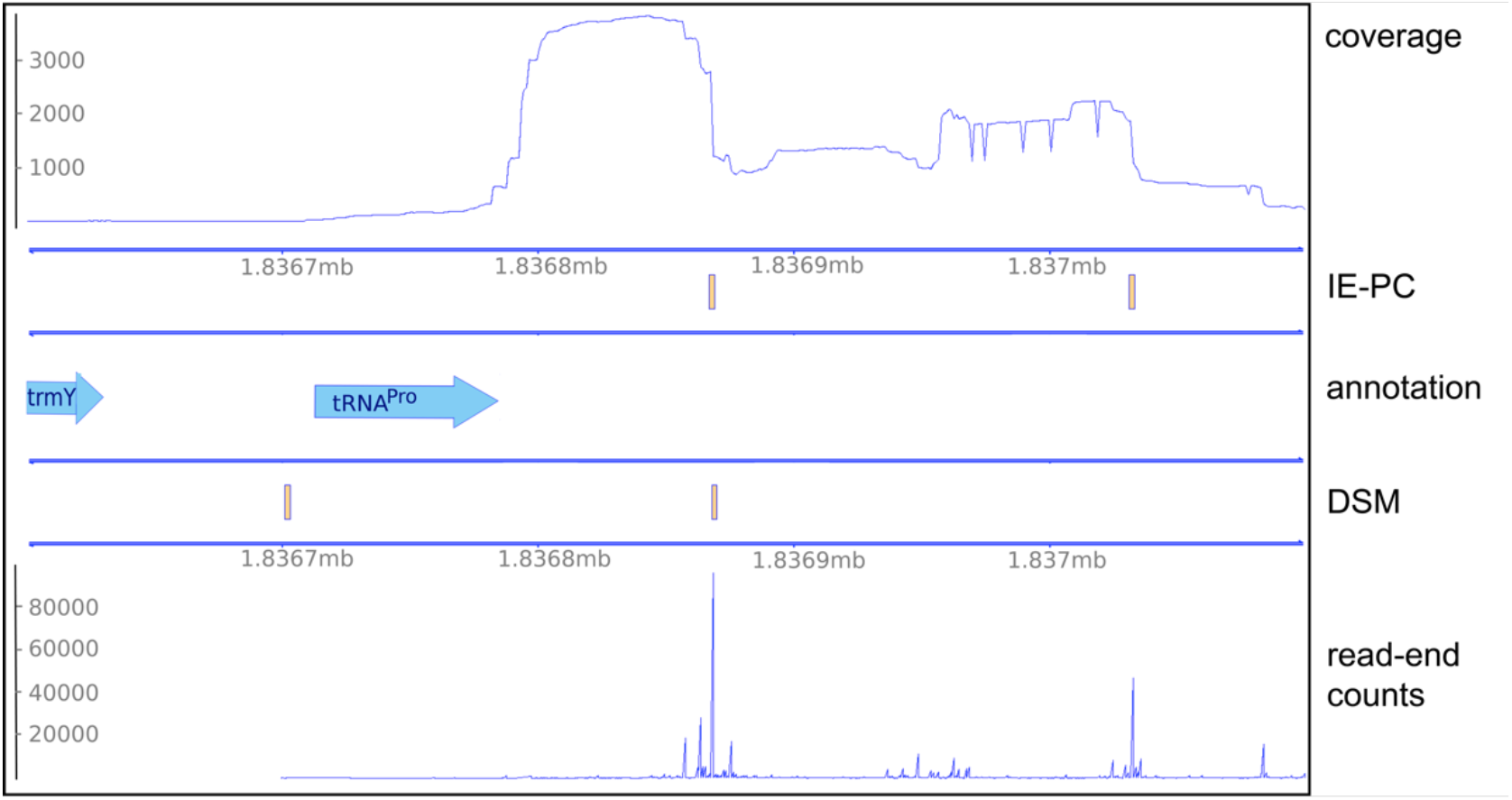
TTS comparison of DSM and IE-PC for the tRNA^Pro^ gene. In the lower panel, termination sites and corresponding read end coverages from the DSM data set are shown, assigning a TTS downstream of the *trmY* gene and another one downstream of the tRNA^Pro^ gene (shown as orange rectangles). The upper panel shows the TTS location determined by IE-PC and the total coverages, corresponding to the -TEX data set, identifying two TTS downstream of the tRNA^Pro^ gene. Since sequencing starts at the 3’ end, coverage starts at the 3’ end and runs continuously for 75 bp due to the read length. The secondary TTS of the tRNA^Pro^ gene was not reported by the DSM algorithm as this algorithm systematically reports only a single TTS for each annotated gene.

**Figure 6.**
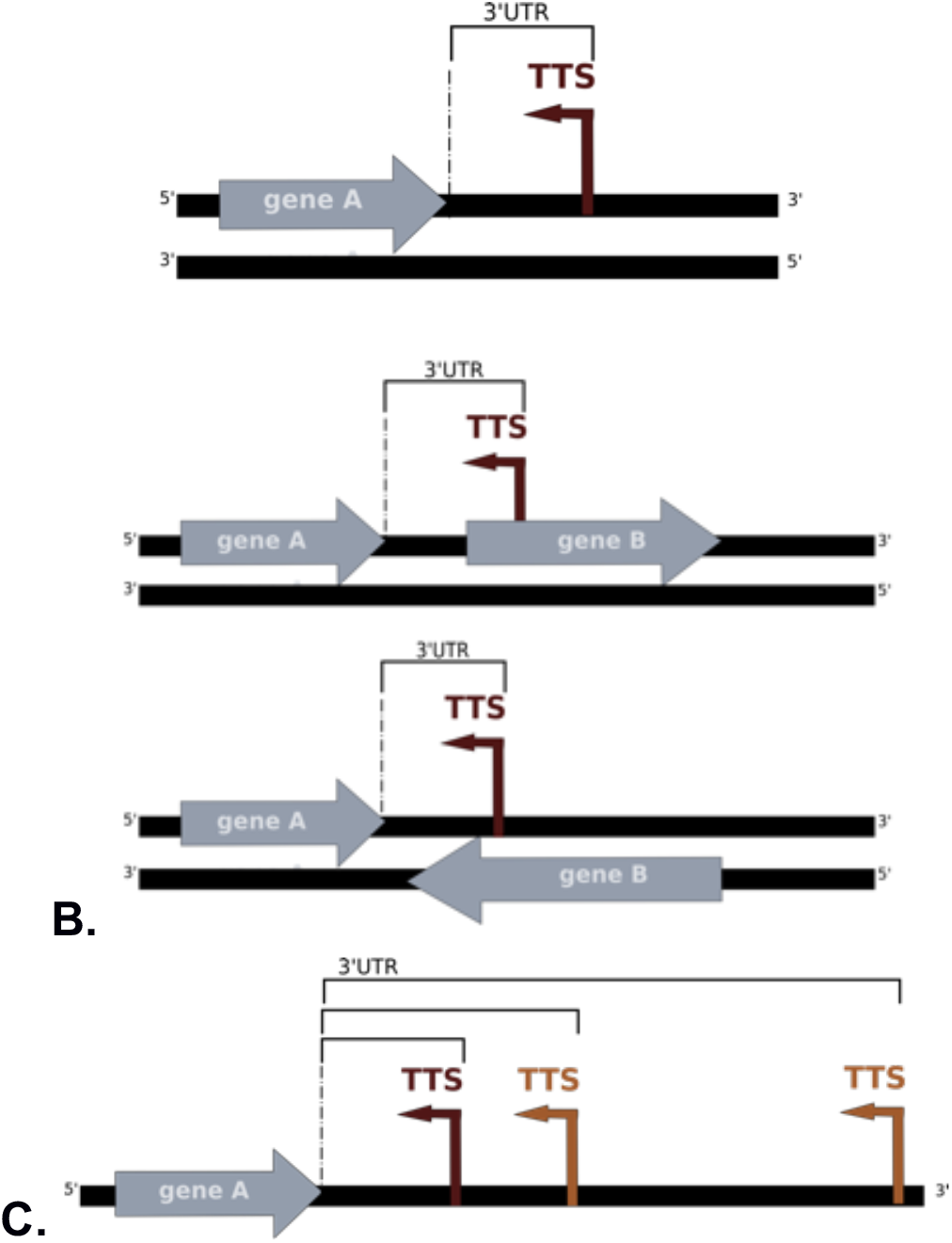
Location of TTS. **A.** TTS is located in an intergenic region. **B.** Location of the TTS in an annotated gene that is located on the sense or antisense strand. The length of the UTR was in all cases measured as the distance between the TTS and the 3’ end of the upstream annotated gene on the same strand. **C.** A first (dark brown) and two secondary TTS (light brown) are shown.

Comparing the DSM data with IE-PC results as well as with given coverage data, we see that chosen downstream regions in DSM were frequently too short.

This is also clear when comparing the median 3’ UTR length determined by both methods. While DSM found a median 3’ UTR length ^1^ of 58 nucleotides, IE-PC determined it to be 96 nucleotides. A comparison of RNA 3’ ends found with DSM and IE-PC for the HVO_1876s gene is shown as an example (Figure 4).

The lower panel (DSM) in Figure 4 shows the results obtained with DSM, coverage is shown as the 3’ end coverage of corresponding reads, the RNA 3’ end position for the transcript is also shown. The upper panel (IE-PC) shows IE-PC results, with RNA 3’ end location and read coverage above. The drop in coverage values is clearly visible, matching the RNA 3’ end location as identified by IE-PC. Corresponding read end coverage of the DSM data set was present at the same position. However, in the DSM analysis, every gene had a downstream region with an individual length, since for each gene the average insert length of the corresponding reads was taken into account. The length for the downstream region of HVO_1876s, was calculated with only 64 nucleotides, and therefore the high read end coverage was beyond the region selected for analysis. As the algorithm is bound to report an RNA 3’ end (there is always a position with highest coverage in the selected region), DSM reported a false positive in this case. For the genes *trmY* and tRNA^Pro^, both methods identified one identical RNA 3’ end position and two different RNA 3’ ends positions (Figure 5). The IE-PC analysis identified two RNA 3’ ends downstream of the tRNA^Pro^ gene (upper panel). The corresponding coverage values for DSM (lower panel) showed similar signals, but due to its specific algorithm, DSM assigned one RNA 3’ end downstream of the *trmY* gene (HVO_1989) and one downstream of the tRNA^Pro^ gene (lower panel).

Taken together comparison between the DSM approach and the IE-PC algorithm clearly shows that using the IE-PC approach yields improved and more comprehensive data.

### Novel approach for the identification of transcription termination sites

To determine RNA 3’ ends the DSM approach used reads from a total cellular RNA fraction that contains RNA 3’ ends derived from transcription termination as well as 3’ ends derived from processing. Thus the 3’ ends identified by DSM are not all TTS but also processing sites (PS). A similar problem exists for the determination of original transcription 5’ ends, where a well established method for the reliable identification of transcription start sites (TSS) has been developed, termed differential RNAseq (dRNAseq). Here, to enrich primary transcripts with original 5’ ends an RNA sample is treated with terminator exonuclease (+TEX) which removes RNAs with a 5’-monophosphate. Primary transcripts are newly synthesized RNA molecules that have not been processed at their 5’ end and also have a higher probability to contain the original 3’ terminus. The majority of bacterial ribonucleases prefer substrates with 5’-monophosphate ends, some even have a specific sensor domain for the 5’-monophosphate (24), thus primary transcripts are less prone to processing.

Therefore -similar to the approach for start site determination-we used a +TEX library to enrich original termination ends. To that end we treated a cellular RNA fraction with 5’ terminator exonuclease (TEX) to enrich primary transcripts and thereby original termination ends. After cDNA library generation from the TEX treated RNA, NGS was performed, resulting in an average of 40 million reads for each of the three libraries (Supplementary Table 10). Reads obtained were analysed with our newly established algorithm as described below.

### The *Haloferax* genome contains 1,543 transcription termination sites

To identify transcription termination sites we applied the IE-PC algorithm to the data from the TEX treated samples consisting of enriched original transcription termination sites, and identified 1,543 putative TTS^2^ (Table 2). Supplementary Table 1 lists all TTS detected.

**Table 2.**
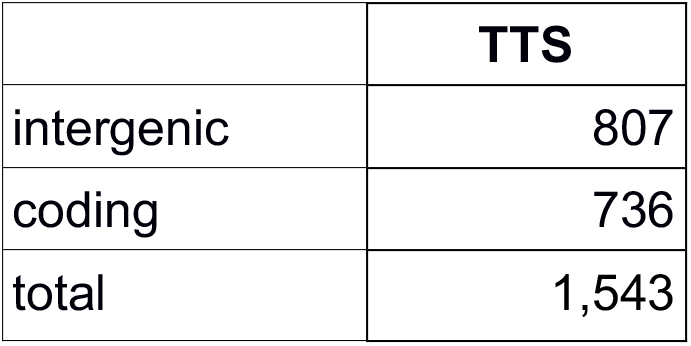
TTS identified with IE-PC. TTS identified are present in coding regions as well as in intergenic regions.

|  | TTS |
| --- | --- |
| intergenic | 807 |
| coding | 736 |
| total | 1,543 |

### Clustering of termination signals

Inspection of the TTS obtained revealed closely spaced TTS, that were 11 to 150 nucleotides apart. These closely spaced TTS were subclassified into first TTS (TTS1) and secondary TTS (TTS_s_): a first TTS is located directly downstream of a 3’ gene end on the same strand (Figure 6A & B); a secondary TTS is located downstream of another TTS, with no other features (like TSS or 3’ gene end) in between, as shown in Figure 6C. We found in total 1,056 first TTS and 487 secondary TTS. We also found very closely spaced TTS that were less than 10 nucleotides apart these were reported as only a single TTS (the one with the highest coverage).

### Distinct motifs for transcription termination in coding and intergenic regions

From the 1,543 transcription termination sites found, slightly more than half of the sites were found in intergenic regions (807 TTS, 52 %) and the remainder in coding regions (736 TTS, 48%)(Table 2). Detailed information for each TTS can be found in Supplementary Table 1.

Analysis of the regions up- and downstream of the TTS were performed separately for TTS located in coding and intergenic regions (Figure 7), and in both, an increase in hybridisation energy at the TTS similar to the increase identified in the TTS set obtained with DSM was found (Supplementary Figure 3). The pattern of nucleotide enrichment for sites located in intergenic regions showed that Ts were prevalent at the TTS (Figure 7A).

**Figure 7.**
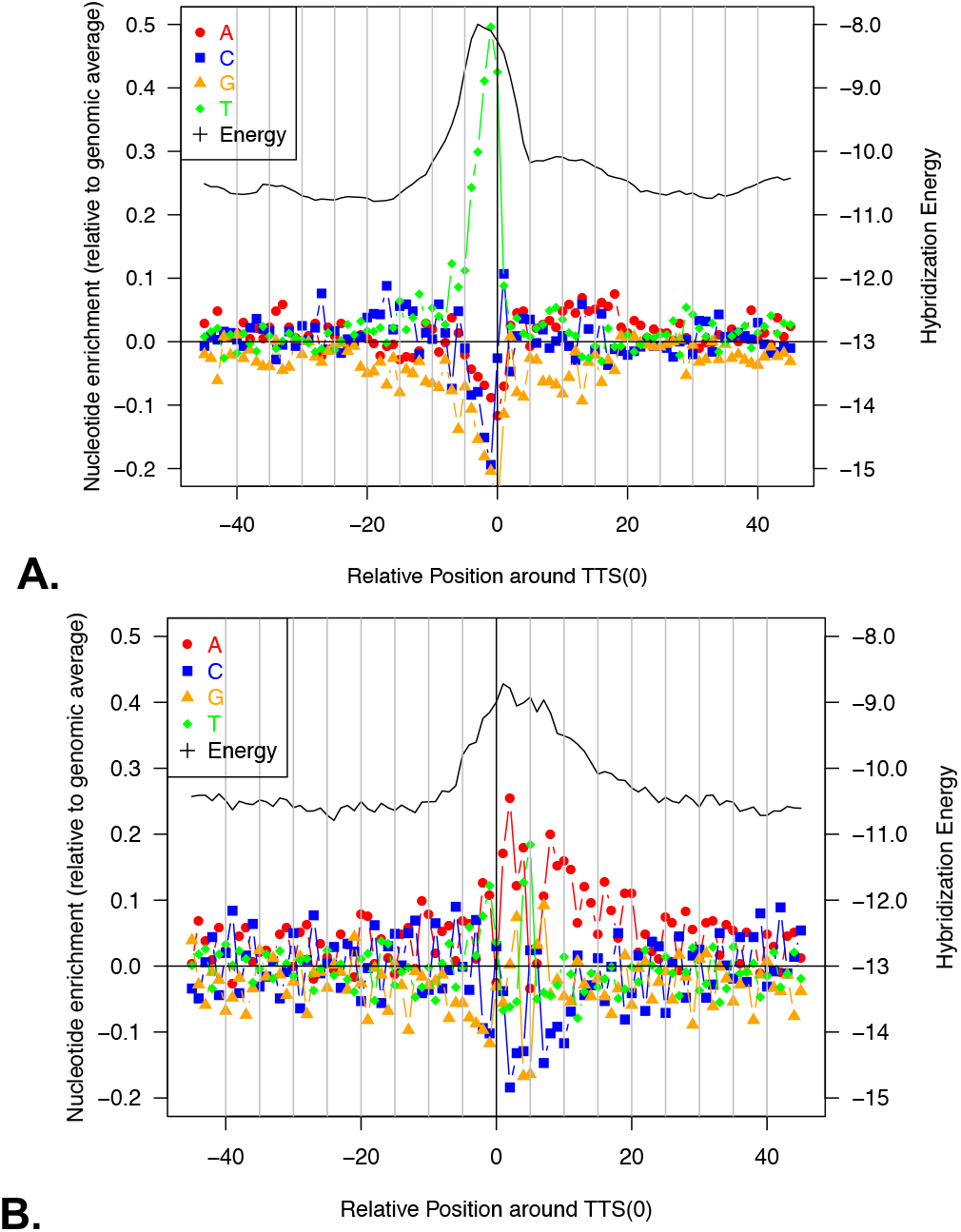
Analysis of up- and downstream regions for TTS identified with IE-PC. **A. Intergenic and B. coding regions were investigated.** Forty-five nucleotides up- and downstream of the termination site were analysed for (1) nucleotide enrichment at each position (left y-axis) and (2) the hybridisation energy (right y-axis). x-axis: nucleotide position (upstream -, downstream +). The colour scheme for the four nucleotides is shown at the upper left, and the energy data are shown with a black line. Hybridisation energies were calculated based on the binding energies between the DNA template and RNA in the area behind the RNA polymerase. The regular pattern in panel B is due to the statistical characteristics of coding regions in GC-rich organisms(25,26).

We next analysed sequences 15 nucleotides up- and five nucleotides downstream of the 807 intergenic termination sites for common sequence motifs. For 748 sites, we found similarities in the sequences, such as a conserved dinucleotide TC as part of the motif as well as a stretch of T’s of variable length upstream of the termination site (Figure 8A, Supplementary Table 2).

**Figure 8.**
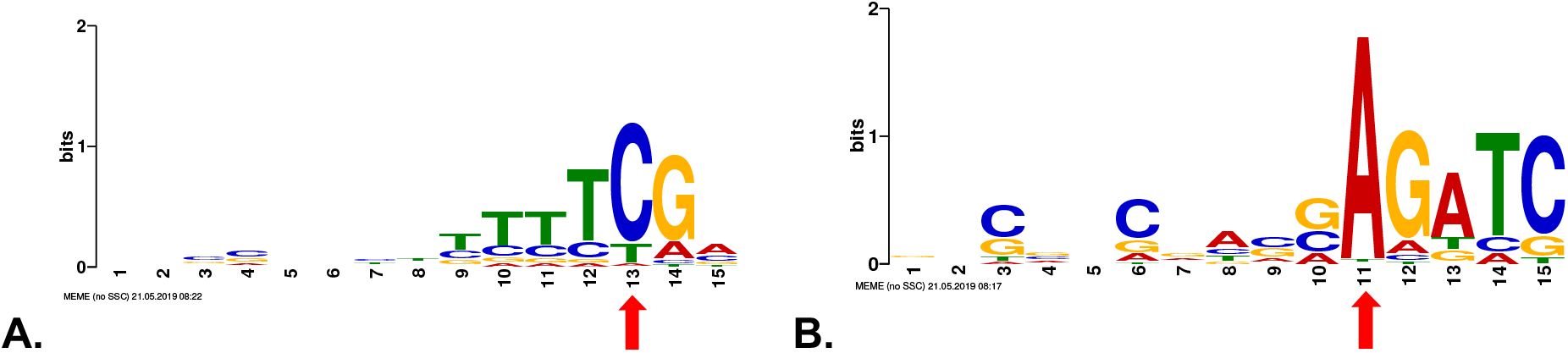
Enriched sequence motifs around the TTS identified with IE-PC. **A.** Sequence motif close to TTS located in intergenic regions. The TTS is located at position 13 (red arrow). **B.** Sequence motif close to TTS located in coding regions. The TTS is located at position 11 (red arrow). Motifs were detected using MEME (27).

Sequences 15 nucleotides up- and five nucleotides downstream of the 736 TTS in coding regions were likewise investigated for common sequence motifs (Figure 8B). The prominent C residue at every third position is typical for coding regions in *Haloferax*. The third codon position combines an enrichment for GC (due to the GC-rich genome) and pyrimidines. For 456 TTS in coding regions, we found the downstream motif AGATC (Figure 8B).

Taken together, we could identify distinct termination motifs that were specific for intergenic and for coding regions.

### Secondary structures can act as termination signals

To identify potential secondary structure motifs, we conducted a search for accessible and inaccessible regions around the TTS using RNAplfold (28) (Vienna RNApackage (29,30)). We plotted accessibilities against a background distribution of shuffled dinucleotides. However, no clear signals were found to indicate significantly increased or decreased accessibility. In a second attempt to identify secondary structures, we applied graphclust 2.0 (31) within Galaxy (32) to sequences 100 nucleotides upstream of all TTS (Supplementary Figure 5). Graphclust is a tool that clusters input sequences based on their secondary structure(s). It will cut the sequences based on a window size parameter and align and fold the sequences into secondary structures using RNAalifold of the ViennaRNA package (29,30). Using this approach we found hairpin structures upstream of 503 of the TTS (up to 10 nucleotides distance), an example is shown in Figure 9.

**Figure 9.**
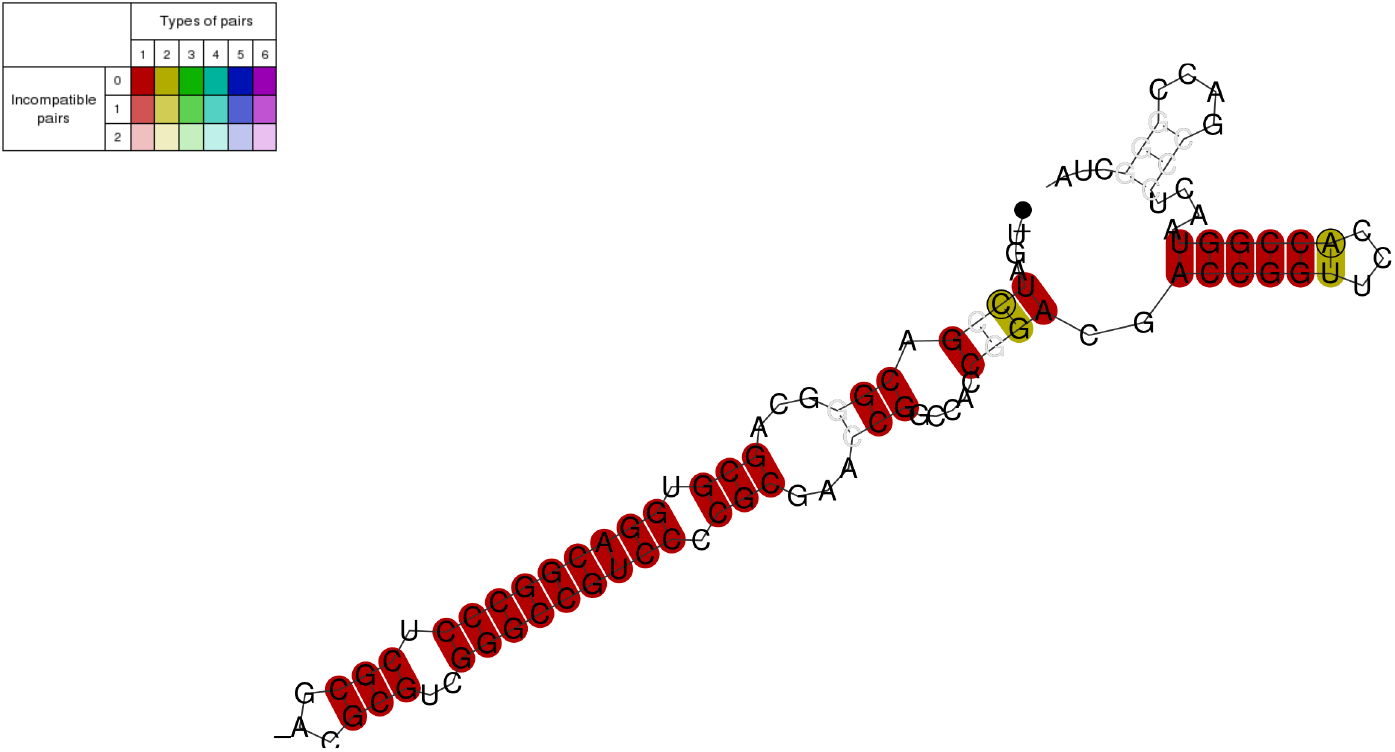
Hairpin structure found upstream of TTS. The secondary structure was plotted with RNAalifold, and it’s 5’ end is denoted by a small dot at the end of the line. The colour coding is at the top left; whereas darker colours show compatible pairs, and the number of types of pairs shows how many types are found at this position thereby indicating sequence conservation. Dark red colours indicate base pairs that are compatible and conserved.

Thus, in some cases secondary structures are present that might influence transcription termination.

### Experimental confirmation of selected termination signals

We selected four TTS identified with our new IE-PC approach in intergenic and coding regions to test their termination activity with an *in vivo* reporter gene assay. Termination regions (the TTS and up- and downstream regions) were cloned into a fragment of the reporter gene β-galactosidase (sequences are listed in Supplementary Table 9). Termination activity was monitored with northern blot analyses (Figure 10, Supplementary Table 13). If termination occurred in the inserted fragment, the RNA was shorter than that in the control construct (Figure 10). The T stretch identified here and in earlier studies with archaea terminated efficiently at the expected site with 95% termination and some read-through, showing that this assay worked well (Figure 10B, Supplementary Table 13). Next, we tested two TTS (A1 and A2) found in coding regions with the downstream motif AGATC. Both terminated at the expected site, confirming the activity of this newly identified downstream motif. However, termination is not as efficient as for the T stretch motif (11% for A1 and 27% for A2) and thus allows considerable read-through (Figure 10C, Supplementary Table 13). Furthermore, a hairpin motif found upstream of TTS in coding regions (S12, shown in Figure 9) was tested in the assay (Figure 10D, Supplementary Table 13). Again, termination was detected at the expected site, 59% of the transcripts are terminated at the structural motif. To exclude that RNAs were processed at the sequence or structure motifs, northern blots were hybridised with a probe against the downstream fragment. If transcription would not terminate at the insert sequence but promote to the final terminator present in the vector and the transcript would be subsequently processed, the downstream processing product would be picked up with this hybridisation. However, no additional RNA fragments were detected (Supplementary Figure 8).

**Figure 10.**
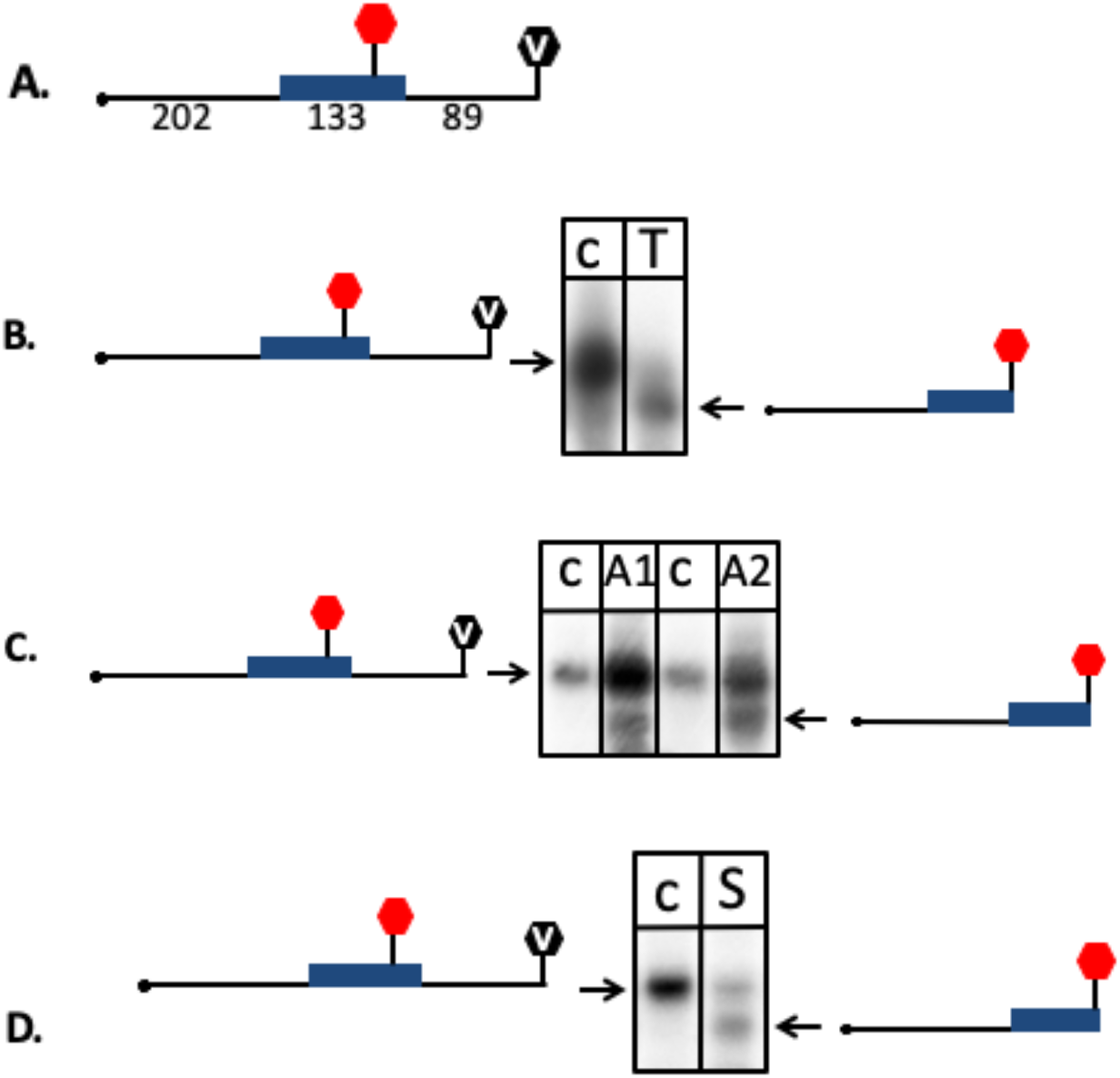
*In vivo* reporter gene test of termination sites. Termination sites were cloned into a reporter gene (the gene for β-galactosidase) construct to confirm their termination activity. RNA was isolated from strains transformed with reporter gene plasmids, separated by size and subsequently transferred to a nylon membrane. Membranes were hybridised with a probe against the β-galactosidase mRNA. Active termination sites generated shorter RNA molecules than the control sequences, indicated by an arrow. Experiments were carried out in triplicate. **A.** The construct used is shown schematically. A full length transcript terminating at the vector termination site is 424 nucleotides long. The insert containing the termination site is 133 nucleotides long (the insert is shown in blue, the location of the termination site in the insert is indicated with a red hexagon), the transcribed plasmid sequences are 202 nucleotides (upstream of the insert) and 89 nucleotides (downstream of the insert) long. The termination site in the plasmid is indicated with a with a V in the black hexagonal. **B.** The T_5_C termination sequence was inserted into the reporter gene. Such a T_5_C sequence can be found for example in the intergenic sequence downstream of the *ftsz2* gene. Transcripts terminating at the T_5_C motif are 321 nucleotides long. The T stretch terminated efficiently (indicated with an arrow) with 95% termination at the T stretch and 5% termination at the plasmid encoded terminator (lane T), lane co: control insert without a termination site. **C.** Two sequences with the downstream motif AGATC (A1 and A2) were inserted into the reporter gene. Both are located in coding regions; A1 is located in the gene HVO_2724, A2 is located in the gene HVO_0455. Termination at these motifs results in 309 nucleotide long RNAs. Both sequences terminated, however considerable read-through is present (termination at A1: 11% and at A2: 27%). Lane A1: AGATC termination site A1, lane A2: AGATC termination site A2. **D.** A sequence with the potential to fold into a hairpin structure (S12) was inserted into the reporter gene. The structure S12 is located in a coding region (gene HVO_A0420). Termination at this motif results in a 309 nucleotide long RNA. S12 terminated, but considerable read-through was observed (termination at S12: 59%). Lane S: sequence S12.

### Analysis of 3’ UTR length

To determine 3’ UTR lengths, the distance between the first TTS and the 3’ end of the preceding annotated gene was determined. The first TTS had a median 3’ UTR length of 97 nucleotides (Figure 11). The median 3’ UTR length for all TTS (first and secondary) is 189 nucleotides. To confirm that the 3’ UTR regions were part of the same transcripts as the upstream coding regions, we generated transcriptome data. To that end we performed next-generation sequencing of a cDNA library generated from total RNA for RNAseq. An average of 42 million reads were obtained for three independent cDNA libraries, and with these data, we confirmed that the 3’ UTR regions were part of the same transcripts as the upstream coding regions, since we found continuous RNAseq reads over the complete putative 3’ UTRs (Supplementary Table 4).

**Figure 11.**
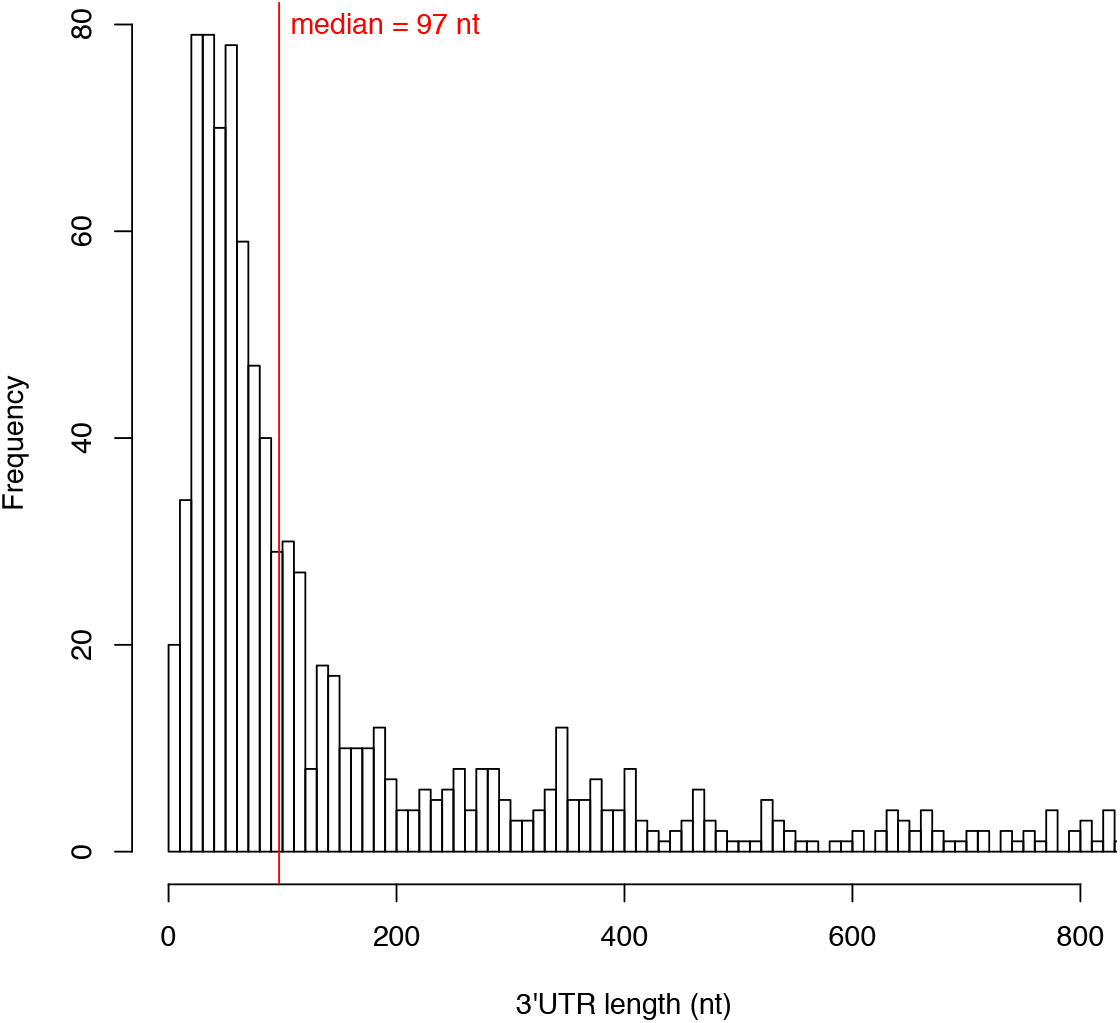
Histogram of 3’ UTR lengths. The 3’ UTR lengths for all first TTS are shown as calculated from the IE-PC data on the basis of the +TEX reads. The histogram shows the frequency of 3’ UTR lengths for the region up to 800 nucleotides in length.

## Discussion

### Novel IE-PC approach allows comprehensive, genome-wide identification of transcription termination sites

Applying the new IE-PC algorithm on +TEX RNA sequences, we were able to determine the TTS genome-wide for the archaeon *H. volcanii*. The dRNAseq method using a +TEX library is an established method to confidently identify transcription start sites by enriching primary transcripts. However, treatement with TEX does not completely remove 5’-monophosphate RNAs, thus only an enrichment of primary transcripts is achieved and the +TEX data set contains to some extent still 5’ processed RNA. Nevertheless, the method is standard and accepted for TSS identification.

Currently very little is known about archaeal RNases and their preferred substrates, but bacterial ribonucleases clearly prefer substrates with a 5’-monophosphate. Thus an RNA with an 5’-triphosphate is not the preferred substrate for these RNases. Therefore, we used here a +TEX library to enrich transcription termination sites. Altogether, we found 1,543 TTS in the +TEX data set with our new algorithm, representing the first unbiased, truly genome-wide approach.

Comparison of both methods for TTS identification, IE-PC and DSM, showed that IE-PC could identify TTS genome-wide, whereas DSM was restricted to regions downstream of annotated genes. Furthermore, algorithmic details of DSM (e.g. the requirement that each annotated gene is directly followed by a TTS; UTR length restriction by using average insert length) caused false TTS identifications in some cases.

### Motifs for termination

Amongst all TTS, two general sequence motifs for termination were identified. One was predominant in intergenic regions, and the other one in coding regions. Both types of termination motifs (T stretch and AGATC) were confirmed using *in vivo* assays. Motifs in intergenic regions consisted of a stretch of Ts, while termination occurred at a C residue. This motif is similar to those found by *in vitro* studies in other archaea (10,12–14,36) and for *S. acidocaldarius* and *M. mazei* using the DSM algorithm (17).

### A specific motif for coding regions

The presence of TTS in coding regions, terminating transcription of an upstream located gene, has been observed previously in archaea including the RNASeq analysis by Dar et al. (17,37). The *in vivo* termination test confirmed that AGATC is used as termination motif but revealed that it is not very efficient (11-27%). This lower termination efficiency is important for TTS in coding regions since it allows full length transcription of the gene in which the termination site is located. A termination event as strong as T_5_C would be detrimental for the expression of the terminator containing gene. In addition, it would be very difficult to place such a T stretch in a coding region since the coding region is governed by the requirement to contain the specific triplets for the encoded protein, and especially a G/C rich genome like the one from *Haloferax* can rarely contain a T stretch in a coding region. Thus, to be able to terminate in coding regions other signals have to be used and these motifs should allow for sufficient read-through.

The prevailing motif we have found in coding regions was AGATC, located downstream of the termination site. The DNA sequence downstream of the TTS has been shown to influence termination efficiency in bacteria and eukaryotes (13). Furthermore, it has been reported that the archaeal RNA polymerase interacts with downstream duplex DNA (38). Thus, it is entirely possible that a downstream termination motif exists. It might also be possible that a protein binds to the downstream motif, acting as a minor roadblock for the RNA polymerase and thereby inducing pausing and in some cases subsequently terminating transcription. Future research will show the mechanistic details of this process.

### Secondary structures involved in termination

The search for secondary structure motifs located close to the TTS was successful for a subset of the TTS and revealed hairpin structures upstream of the TTS, suggesting that these structures might influence transcription termination. One of the identified structures was tested in the *in vivo* termination experiments and indeed terminated transcription, however not as efficiently as the T stretch. A certain amount of folding potential upstream of TTS was also found in *M. mazei* but was considered not to be essential, whereas no structures were identified near TTS in *S. acidocaldarius* (17). Possibly, termination signals in different archaeal phyla are more variable.

### Different 3’ UTR lengths are generated by termination at several TTS

With our genome-wide TTS determination approach, we can present the first comprehensive determination of 3’ UTR lengths in *H. volcanii*. The median 3’ UTR length for first TTS was 97 nucleotides, which is longer than the median 3’ UTR length recently reported for two archaea (with 55 nucleotides for *S. acidocaldarius* and 85 nucleotides for *M. mazei*) and the bacterium *B. subtilis* (40 nucleotides)(17). The presence of transcripts with long UTRs was confirmed by RNAseq data.

In many cases, we found several TTS downstream of one gene, showing that mRNAs with different 3’ UTR lengths are generated. The lengths of the UTRs as well as the fact that some genes have several 3’ UTRs provide a good platform for interactions with regulatory molecules, such as sRNAs. This is similar to the situation in eukaryotes, in which termination at different sites generates different 3’ UTRs, that can interact with different regulatory proteins or RNAs such as miRNAs. Different UTR lengths were also found in *M. mazei* and *S. acidocaldarius* (17).

### Transcription produces antisense RNA

If a gene is located on the opposite strand of a TTS (Figure 6B, lower panel), an antisense RNA against this oppositely encoded gene is generated. Of the 1,543 TTS detected, 14.4% (222 TTS) of them were located in a gene on the opposite strand, thus resulting in antisense RNAs. Up to 75% of all genes were found to be associated with antisense RNAs in bacteria (39–41). Studies with archaea painted a similar picture (21,42). In *S. acidocaldarius*, 301 gene pairs with convergent orientations were investigated, and in 52% of them, the terminator of one gene was located in the coding region of a gene on the opposite strand (17) (as in Figure 6B lower panel); however, in *M. mazei*, only 8% of the convergent genes showed such an overlap. The potential functions and the physiological relevance of these antisense transcripts must be uncovered in the future.

### Conclusion

Taken together, we showed that our new IE-PC approach used with a +TEX data set is well-suited to identifying the complete set of TTS for a genome without any a priori limitations on the search space due to genome annotation. We confirmed the T stretch termination motif detected 30 years ago, but we also identified an additional new motif specific for coding regions as well as a hairpin structure for termination. The presence of multiple 3’ UTRs for a gene provides a platform for regulatory mechanisms similar to those described in eukaryotic systems.

## Materials and Methods

### *Haloferax volcanii* culture conditions and RNA extraction

*H. volcanii* strains H119 (Δ*leuB*, Δ*pyrE2*, Δ*trpA*)(43) and HV55 (ΔleuB, ΔpyrE2, ΔtrpA, Δ*bga*H) (this work, see below) were grown aerobically at 45°C in Hv-YPC or Hv-Ca medium (43) to an OD_600_ of 0.8-0.9 (for a list of strains used see Supplementary Table 9). Total RNA was isolated from three biological replicates using TRIzol (ThermoFisher scientific). RNA fractions were sent to vertis (vertis Biotechnologie AG, Martinsried, Germany) for further treatment, cDNA library preparation and high throughput sequencing. RNA preparations were made for each of the three different RNAseq approaches: (1) To obtain an RNA fraction enriched in primary transcripts RNA was treated with terminator exonuclease (this fraction was termed +TEX), (2) RNA from one preparation was left untreated representing the complete RNA pool (this fraction was termed -TEX). RNA preparations (1) and (2) were performed such that the original RNA 3’ end was maintained (for details see Supplementary methods). For RNA preparation (3) RNA was isolated for obtaining transcriptome data (this fraction was termed RNAseq data). In all sequencing approaches primers were designed such that the identification of the RNA strand was possible. Details for library construction and sequencing are reported in Supplementary methods.

### *E. coli* culture conditions

*E. coli* strains DH5α (Invitrogen) and GM121 (44) were grown aerobically at 37°C in 2YT medium.

### Read mapping

Raw reads were adapter clipped and quality trimmed using cutadapt version 1.10 (45) based on fastqc version 0.11.4 (46) quality control reports. Reads were then mapped with segemehl (version 0.2.0) (47,48). We used -A 94 to require higher accuracy in order to account for prokaryote mapping instead of mapping eukaryotic genomes. Mapped reads were afterwards processed using samtools version 1.3 (49). In order to calculate genome coverage and intersection of data sets, we used bedtools (bedtools v2.26.0)(50).

### DSM analysis

Based on the Dar-Sorek-Method (DSM)(17,51), RNA 3’ ends (putative TTS) were retrieved from the mapping data. We used samples without TEX treatment for consistency with the DSM analyses and we only used read pairs and at least a coverage of 4 for every valid position. As described in Dar et al. (17,51), for each annotated region in the genome, all inserts overlapping with the annotated region were collected. Then, for each position downstream of a gene, all collected reads that end at this position were summarised, excluding inserts longer than 500 bp. The average insert length of all the corresponding read pairs was defined as the length of the target downstream region. For all read pairs where the insert overlapped an annotated region, the position with highest read-end coverage inside the target downstream region of an annotated 3’ end was reported as a transcription termination site (TTS) (Table 1 and Figure 1). Collected TTS with DSM are listed in Supplementary Table 11. The used implementation of this method is available at Bioinformatics Leipzig (http://www.bioinf.uni-leipzig.de/publications/supplements/18-059).

### Internal Enrichment algorithm

To detect sites with a significant enrichment of sequenced and mapped fragment ends a sound background without enrichment is desired. In the current setting, we used the intrinsic properties of a paired-end sequencing run to directly deduce the following information (Figure 2). We also took advantage of the fact, that the cDNA library preparation applies a fragmentation step after ligation of the 3’ end primer. Since each fragment which results from an individual fragmentation event (in contrast to PCR duplicated fragments) is very unlikely to have the exact same length, truly enriched sites can be expected to be associated with sequenced fragments, all ending at the respective site but starting at different positions. Therefore, the more different mates (mapping to different positions) are associated with the different reads ending at a particular site the higher the enrichment of read end signal at that particular position can be considered. To capture this, we calculate for each position i a score S as

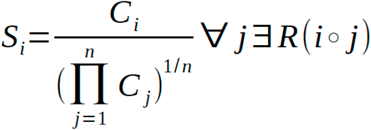

Thereby, C_i_ denotes the number of fragment ends at position i, C_j_ the number of fragment starts at position j, and *R*(*i* ∘ *j*) all position i,j which are associated via at least one read-mate-pair R. To get an expected background distribution of these scores, we again use the nature of paired-end reads. Since we expect only fragment ends, in contrast to fragment starts, to be enriched, we can use the distribution of the reciprocally defined scores for the fragment start S_j_ as a background distribution. Based on the distributions, native fragment end scores can be assigned an empirical p-value, evaluating its likelihood to occur by chance alone. The above detailed software, named Internal-Enrichment (IE) was implemented in Perl and is available at Bioinformatics Leipzig (http://www.bioinf.uni-leipzig.de/publications/supplements/18-059). We ran IE using all position showing signals for starts and ends (-mr 1), we omitted read fragments with length more than 100 nt (-mf 100) to reduce outliers, and the geometric mean as a method to calculate a score for the given signals (-mode GeomMean). We only included signals that were present in all three replica such that each showed a maximal empirical p-value of 0.05. The geometric mean of all empirical p-values for one position from all replica is used as the final score for the given position.

### Peak Calling

A complementary approach to find RNA 3’ ends is identifying peaks in read stops (Figure 3). In order to find these peaks, we first computed the strand specific read coverage at every position in the genome, which we used as a background. We then used a sliding window approach with a window size of 150 nt and an overlap of 50 nt to find positions in the respective windows with a significantly higher number of read stops than the rest of the window. This was done by computing the mean number of stops as well as the standard deviation by window and then calculating the z-score for the number of read ends at every position of the window. Subsequently, we required a minimum number of 5 reads stopping, at least 10% of all reads covering position i-1 must end at position i as well as a minimum z-score of +2 at a site to report it as a putative RNA 3’ end. As a consequence of the overlapping of the windows, a site is evaluated multiple times and must fulfil the criteria in at least one contexts evaluated.

### Enrichment of primary transcripts

To be able to differentiate between 3’ end processing sites and termination sites we isolated RNA and enriched this RNA for primary transcripts by digestion with the terminator exonuclease (TEX) (for details see Supplementary methods). Primary transcripts contain 5’ triphosphates and are not digested by TEX. Bacterial ribonucleases preferential use 5’-monophosphate RNA as substrates (24), thus the RNA fraction enriched for 5-triphosphate RNAs should be enriched in unprocessed RNAs.

### Terminator identification

We report results for the sample enriched in primary transcripts (+TEX), in which IE and PC were both computed for the +TEX sample, followed by intersection analysis, resulting in the identification of TTS. In most cases, both methods (IE and PC) reported the same sites. For a few cases, more than one position was reported within a distance of 10 nt. In this case, the position with the highest coverage was chosen as TTS. More information is available at Bioinformatics Leipzig (http://www.bioinf.uni-leipzig.de/publications/supplements/18-059).

### Terminator sequence and structure analysis

We calculated percentages of nucleotide enrichment and hybridization energies for regions of 45 nt upstream and downstream of the TTS. Hybridization energies are calculated as the energy that stabilizes the RNA-DNA hybrid during transcription (28), using RNAplfold, a program part of the ViennaRNA package (30) (version 2.4.9) with special energy parameters for RNA-DNA hybrids. The lower (the more negative) the hybridization energy, the more stable is the RNA-DNA hybrid.

### Motif and structure analysis for 3’ UTR sequences

Motifs were detected using MEME (27) (version 5.0.1) by scanning all sequences from 15 nt upstream to 5 nt downstream of the TTS (see Figure 8 for an example). Structure search was conducted using graphclust 2.0 (31) within Galaxy (32). Input sequences were sequences of 100 nt length upstream to the TTS. Graphclust was used with default parameter and additionally a window size of 110 (such that our sequences fit in one window to avoid duplicated sequences), a bitscore of 15 for the results of cmscan, an upper threshold of 50 clusters and 20 top sequences in each alignment for the visualization. The application of graphclust resulted in 37 clusters providing covariance models (CMs) for each. Covariance models are probabilistic models that are created based on given sequence and structure motifs and can be used to search for sequence and structure homologies. To scan sequences for these given motifs, we applied cmscan (a part of the infernal program suite (52), version 1.1.1) on our input sequences after calibrating the CMs (using cmcalibrate, part of the infernal program suite) and additionally on all the transcripts in order to get a background model. Given cmscan results, we filtered the secondary structures detected in the TTS-related sequences such that the structures were at most 10 nt upstream from the TTS.

### Analysis of 3’ UTR regions

We confirmed 3’ UTR regions by checking if coding region and 3’ UTR are completely covered by RNAseq reads. If so, this would be indicative of uninterrupted transcription until the assigned TTS. Out of 1,543 3’ UTR sequences, 1,286 show a continuous coverage within RNAseq reads (82.2%) (Supplementary Table 4).

### Code availability

The code is available at: http://www.bioinf.uni-leipzig.de/publications/supplements/18-059 (see also data availability statement below).

### Reporter gene investigations of identified TTS

*Construction of bgaH deletion mutant HV55.* To allow the use of a plasmid carrying the ß-galactosidase reporter gene (a fusion of ß-galactosidase genes from *Haloferax alicante* and *Haloferax volcanii*)(*bga*Ha)(53,54), the *H. volcanii* ß-galactosidase gene (*bga*H) was deleted in strain H119. H119 was transformed with the *bga*H deletion plasmid pTA617 (for a list of plasmids used see Supplementary Table 9)(55,56) and grown in Hv-Ca medium with tryptophan (final concentration 0.25 mM) to generate the deletion strain (57). Homozygous knock-out clones were verified via southern-blot using *Sal*I digested genomic DNA and probe bgaHaDO (primers: bgaHKODO-for and bgaHKODO-rev (for a list of primers used see Supplementary Table 9); template genomic DNA of *H. volcanii*). Probe labelling and detection of the blot were carried out using the DIG-DNA labelling mix and detection reagents (Anti-Digoxigenin-AP) (Roche) according to the manufacturer’s protocol. The resulting deletion strain was termed HV55.

*Construction of pTA231-termtest constructs.* To construct the terminator-test plasmid a p.*syn* expression cassette (Anice Sabag-Daigle and Charles J. Daniels, in preparation) synthesised by lifetechnologies^TM^ (Thermofisher Scientific) was introduced via *Not*I/*Eco*RI into pTA231 (43), resulting in pTA231psyn (for sequences see Supplementary Table 9). This plasmid was subsequently cured of the *Bam*HI site by digestion with *Bam*HI, blunting by Klenow fragment and religation. Then a C-terminal fragment of the reporter gene *bga*Ha was inserted at the *Nde*I site. After *Nde*I digestion, the vector was treated with Pfu polymerase to fill-in the *Nde*I site. The inserted *bga*Ha fragment was generated by PCR using primers bgaHatermifw and bgaHatermirev and pTA599 as template (55). The correct orientation of the inserted fragment was confirmed by sequencing, the resulting plasmid was termed pTA231-termtest. The candidate terminator sequences were inserted into the *Bam*HI site that is present in the newly inserted fragment. Terminator fragments were generated with PCR using primers as listed in Supplementary Table 9 and templates pTA599 (for the control construct) or genomic DNA of *H. volcanii* (for TTS-A1, TTS-A2, TTS-S12 and TTS-ftsZ). HV55 cells were transformed with pTA231termtest constructs, and grown in Hv-Ca medium with uracil (final concentration 0.45 mM). Cells were grown to an OD_650_ of 0.6-1.0 and RNA was isolated using NucleoZOL (Machery-Nagel) according to manufacturer’s instructions.

*Northern-blot analysis of Hv55xpTA231-termtest RNA.* To analyse the RNA levels, total RNA was isolated from *H. volcanii* as described above. Ten µg RNA was separated on a 1.5% agarose gel and subsequently transferred to a nylon membrane (Biodyne® A, PALL). After UV-crosslinking the membrane was hybridised with a radioactively labelled probe against the 5’-part of the *bga*Ha mRNA to detect termination events. The probe was generated by PCR (primers bgaHatermi fw and TermiVectorrev; template pTA599) and the purified PCR fragment was labelled using α-^32^P-dCTP and the random primed DNA labelling kit DECAprime^TM^II (invitrogen). Experiments were done in triplicate. For detection of a potential processing fragment downstream of the termination site, a radioactively labelled probe against the 3’-part of the *bga*Ha mRNA was used. The probe was generated and labelled as described above using primers BetaHinten1 and BetaHinten2 and pTA231-termtestA1 as template. For quantification of northern blot signals, membranes were exposed to phosphorimaging plates (FujiFilm) and the resulting signals detected using a Typhoon imager (GE). Analysis of three replicates was carried out using the ImageQuant TL software (GE).

### Data availability

The data supporting our findings are available in the Supplementary Information, in addition the code is available under (http://www.bioinf.uni-leipzig.de/publications/supplements/18-059), RNAseq data are available at the European Nucleotide Archive (ENA; https://www.ebi.ac.uk/ena) under the project accession number PRJEB30349, with the following assigned experiment accession numbers.

| Experiment replicon | Experiment accession number at ENA |
| --- | --- |
| + TEX, replicon 1 | ERX3014341 |
| + TEX, replicon 2 | ERX3014376 |
| + TEX, replicon 3 | ERX3014377 |
| - TEX, replicon 1 | ERX3014378 |
| - TEX, replicon 2 | ERX3014379 |
| - TEX, replicon 3 | ERX3014380 |
| transcriptome, replicon 1 | ERX3014381 |
| transcriptome, replicon 2 | ERX3014382 |
| transcriptome, replicon 3 | ERX3014383 |

## Acknowledgements

AM, LKM, JW and PM would like to thank Britta Stoll, Elli Bruckbauer, Irma Merdian and Susanne Schmidt for expert assistance.

## Footnotes

1 Here the distance between the 3’ end of a gene and the TTS found directly downstream is determined.

2 For better understanding RNA 3’ ends identified from the +TEX data are termed TTS throughout the manuscript, all ends identified using the -TEX data are termed RNA 3’ ends.

