## Supplemental Data for "Identification of RNA 3’ ends and termination sites in *Haloferax volcanii*"

**Supplementary Data**

Supplementary Data consist of:

A. Supplementary Methods (page 1),
B. Supplementary Figures 1-6 (page 2),
C. Legends to Supplementary Tables 1-4 (page 7),
D. Supplementary Tables 5-8 (page 9),
E. References (page 13).

### A. Supplementary Methods

Library construction and next generation sequencing

All RNA fractions were treated with RiboZero kit (Epicentre) to remove the ribosomal RNAs. The RNA fraction "+TEX" was first incubated with T4 polynucleotide kinase (to phosphorylate 5´ hydroxyl groups) and subsequently with Terminator exonuclease (TEX) (Epicentre) to remove the RNA species which carry a 5´ mono-phosphate, thus this RNA fractions is enriched in primary transcripts. After the TEX treatment an adenylated oligonucleotide (3´ Illumina sequencing adapter) was ligated to the 3´ OH ends of the RNAs. Then, first-strand cDNA synthesis was performed using M-MLV reverse transcriptase and the 3´ adapter as primer. The first-strand cDNA was fragmented and to the 3´ ends of the cDNA fragments the 5´ Illumina sequencing adapter was ligated. Finally, cDNAs were PCR-amplified using a high fidelity DNA polymerase (Herculase II Fusion DNA Polymerase, Agilent 600679) and TruSeq Dual Index PCR primers.

The RNA fraction "-TEX" was not treated with TEX. An adenylated oligonucleotide (3´ Illumina sequencing adapter) was ligated to the 3´ OH ends of the rRNA depleted RNAs. Then, first-strand cDNA synthesis was performed using M-MLV reverse transcriptase and the 3´ adapter as primer. The first-strand cDNA was fragmented and to the 3´ ends of the cDNA fragments the 5´ Illumina sequencing adapter was ligated. Finally, the cDNA fragments were PCR-amplified using a high fidelity DNA polymerase (Herculase II Fusion DNA Polymerase, Agilent 600679) and TruSeq Dual Index PCR primers.

For preparation of the RNA fraction RNAseq (also termed transcriptome), the RNA was first fragmented using ultrasound (4 pulses of 30 s each at 4°C). Then, an oligonucleotide adapter was ligated to the 3´ end of the RNA molecules. First-strand cDNA synthesis was performed using M-MLV reverse transcriptase and the 3´ adapter as primer. The first-strand cDNA was purified and the 5´ Illumina TruSeq sequencing adapter was ligated to the 3´ end of the antisense cDNA. The resulting cDNA was PCR-amplified to about 10-20 ng/µl using a high fidelity DNA polymerase (Herculase II Fusion DNA Polymerase, Agilent 600679).

For all RNA fractions the cDNAs were purified using the Agencourt AMPure XP kit (Beckman Coulter Genomics) and were analysed by capillary electrophoresis. For Illumina sequencing, the cDNAs were pooled in approximately equimolar amounts. The cDNA pool was eluted in the size range of 200 –500 bp from a preparative agarose gel. An aliquot of the size fractionated cDNA pool was analysed by capillary electrophoresis. The cDNA pool was sequenced on a Illumina HiSeq 2000. The number of obtained reads, mapped and unmapped reads, respectively, is shown in Supplementary Table 10.

**B. Supplementary Figures 1 - 6**

**Supplementary Figure 1. 3´ UTR lengths identified with DSM.** Histogram showing the length distribution of the identified 3´ UTRs. The median 3´ UTR length was 58 nucleotides. x-axis: 3´ UTR length (distance between TTS and 3´ end of annotated upstream gene), y-axis: frequency of data points falling inside one of the equi-distanced areas of the histogram.

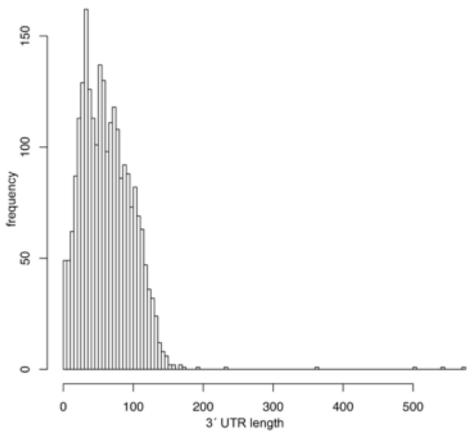

**Supplementary Figure 2. Termination site for the *pilA2* gene.** The termination site downstream of the *pilA2* gene (HVO_2062) is shown. The termination site is indicated by an orange rectangle in the middle panel. The annotated gene is shown in the upper panel, and the genomic location is indicated below the annotation and above the coverage as a blue line with coordinates given in Mb. The read end counts are shown in the lower panel (read counts per position), and DSM data are reported as binary signals; thus, either a signal is present or not. The Figure was created using the R package Gviz[23].

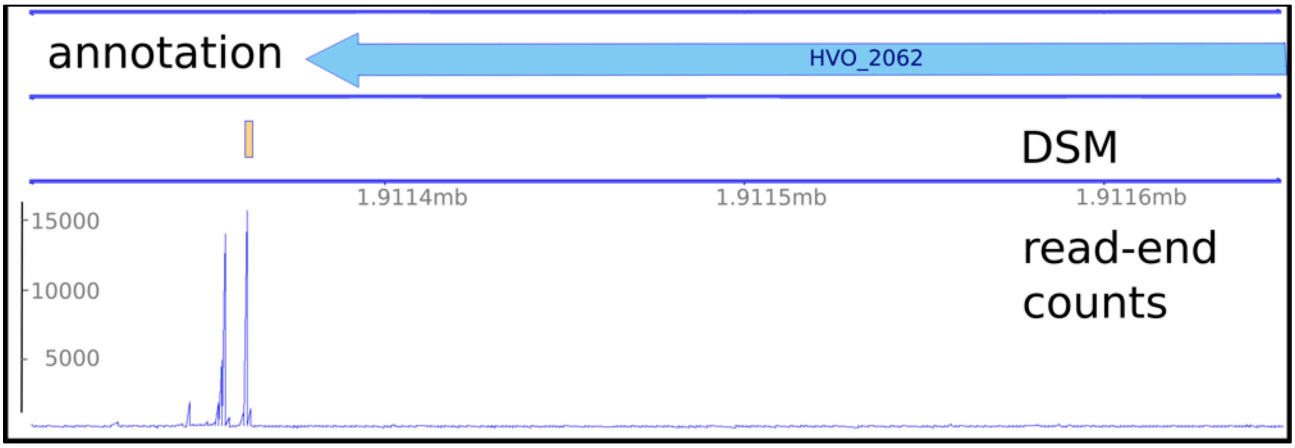

**Supplementary Figure 3. Analysis of up- and downstream regions for TT sites identified with DSM. A. Intergenic and B. coding regions were investigated.** For the TTS identified with DSM, 45 nucleotides up- and downstream of the termination site (x-axis) were analysed in terms of a) nucleotide enrichment at each position (left y-axis) and b) the hybridisation energy (right y-axis). The colour scheme for the four nucleotides is shown at the upper left, and the energy data are shown with a black line. Hybridisation energies were calculated based on the binding energies between the DNA template and RNA in the area behind the RNA polymerase (for details, see Materials and Methods).
Determination of the hybridisation energy ∆G between the nascent mRNA and the template DNA around the TTS revealed that the hybridisation energy increased at termination sites, which indicates a destabilisation of the DNA-RNA hybrid, supporting transcription termination. This was especially prominent for sites in intergenic regions. Nucleotide enrichment analyses for the intergenic TTS showed a clear preference for thymidine two to seven nucleotides upstream of the TTS. The number of adenosines was slightly higher five to 21 nucleotides downstream of the TTS. The region upstream of the TTS (up to position -8) was depleted of cytidine (with the strongest C depletion at position -3) and guanidine nucleotides (with the strongest G depletion at -1). The common motif for termination, as deduced from the nucleotide enrichment, showed that T frequently occurred directly upstream of the termination site, with C higher than average close to the termination site, A being enriched downstream of the termination site and G being depleted. Within coding regions, T enrichment upstream of the termination site and A enrichment downstream were found. C and G were both depleted around the TTS.

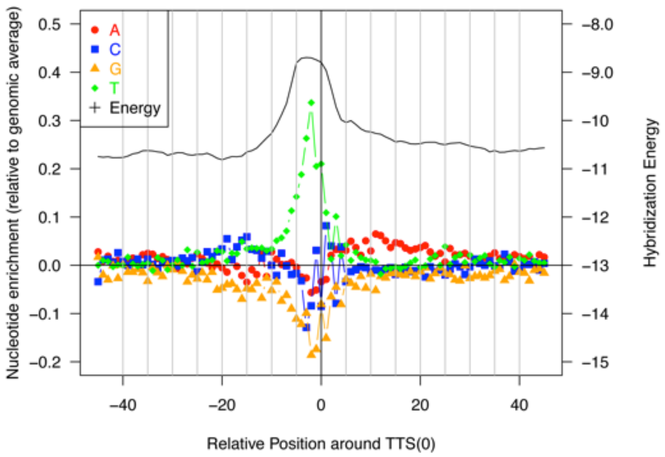

**A.**

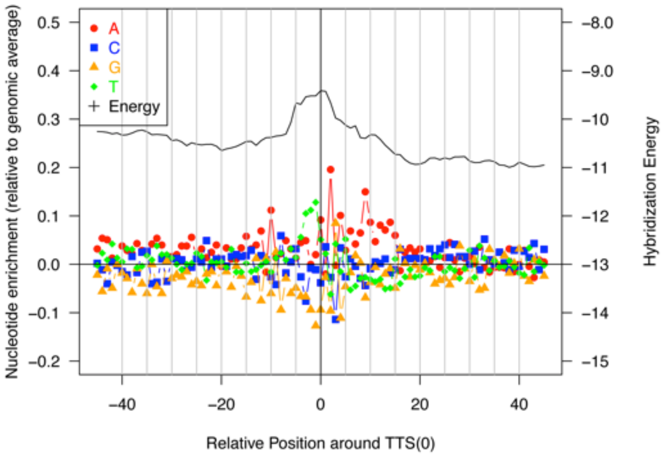

**B.**

**Supplementary Figure 4. Alignment of sequences upstream of the TTS.** The alignment is part of the graphclust output and shows the 20 sequences that fit best (compare also Figure 9). Sequence identifiers are internal graph clust identifiers. The TTS is located at position 100, as sequences were cut with 100 nt upstream and 2 nt downstream of the TTS.

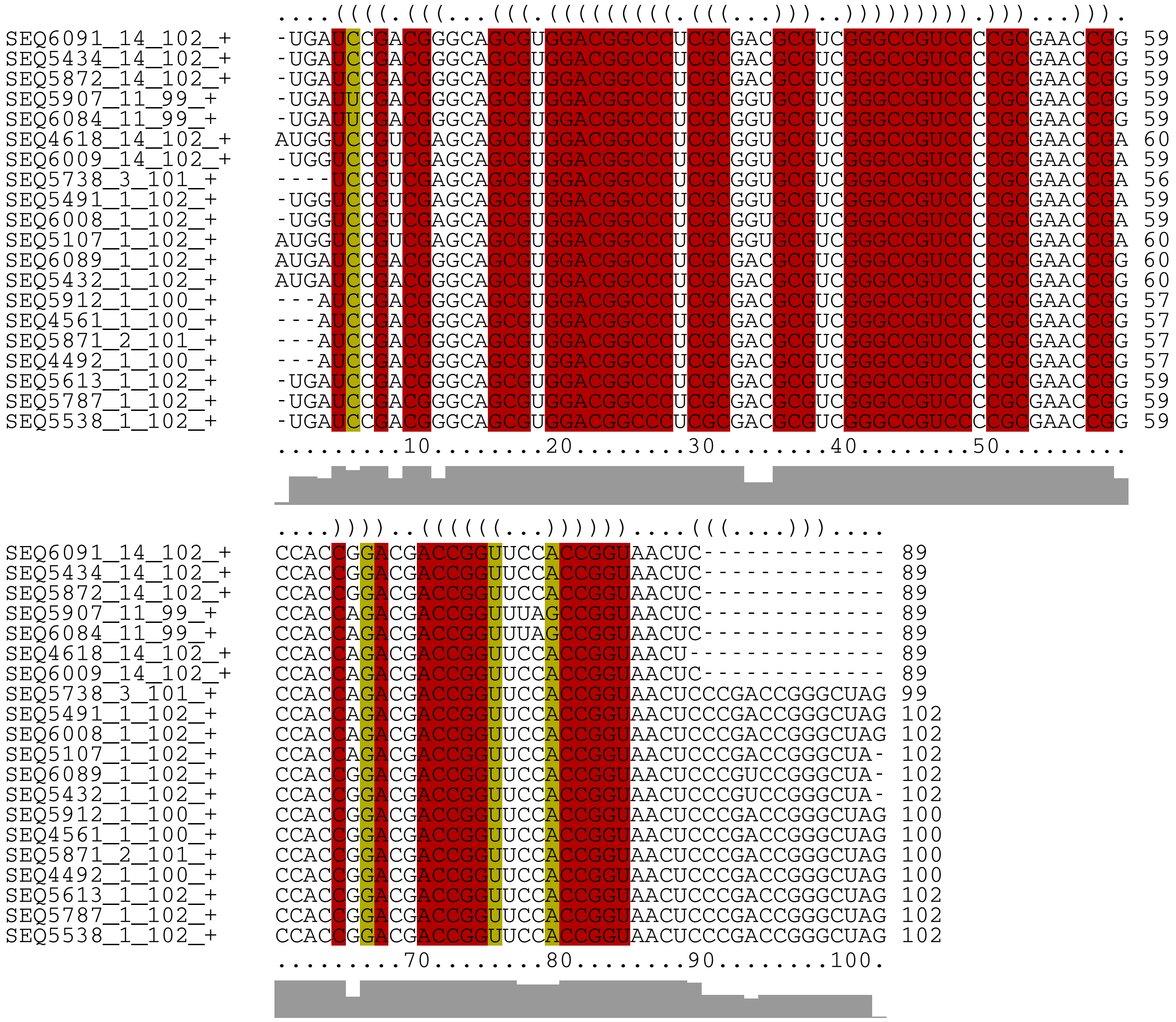

**Supplementary Figure 5. A dicistronic operon with separate promoters and terminators for each gene.** The dicistronic operon encodes the Lsm (HVO_2723) and the Rpl37e (HVO_2722) proteins. Experimental data showed that both genes are transcribed from the first promoter upstream of the *lsm* gene (green arrow upstream of the *lsm* gene)[40]. The *rpl37e* gene also has an independent promoter located in the *lsm* gene (green arrow therein)[40]. Transcription of the *lsm* gene can terminate at the first terminator downstream of the *lsm* gene (red arrow in *rpl37e* gene). A second terminator downstream of the *rpl37e* gene can terminate the dicistronic *rpl37e/lsm* mRNA or the monocistronic *rpl37e* transcript.

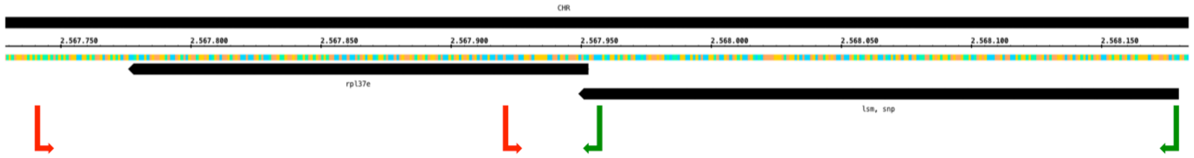

**Supplementary Figure 6. Transcripts are not processed at the AGATC motif.** *Haloferax* cells were transformed with plasmids containing the AGATC motif, RNA was isolated from cells, separated on denaturing agarose gels and transferred to membranes. Membranes were hybridised with a probe against the sequence downstream of the AGATC motif**. A.** Overview over the type of RNAs generated by termination and **B.** by processing. **A.** If AGATC is used for termination, hybridisation with the downstream probe will detect only the full length transcript that terminates at the vector terminator (424 nucleotides long). **B.** If AGATC is a processing signal, the probe should detect the downstream processing product and the full length transcript (depending on how efficient processing is) (424 nucleotides and approximately 100 nucleotides long). **C.** Northern detection. Lane A1 and A2: RNA isolated from cells with the plasmid containing the A1 and A2 sequence, respectively. The probe detects only the full length transcript (424 nucleotides long), that terminates at the vector terminator. Shorter RNA molecules are not detected (processed product: approximately 100 nucleotides long), showing that the RNA is not processed.

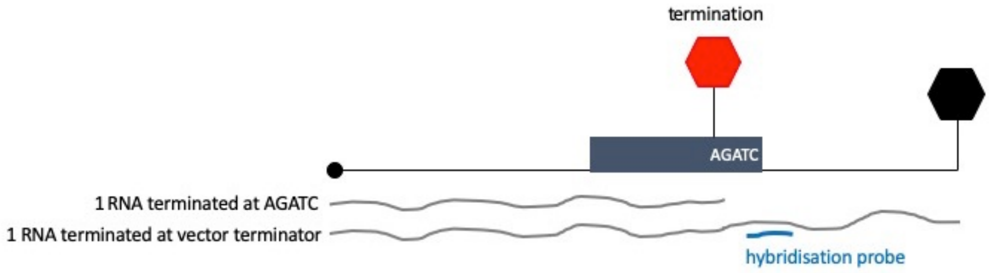

**A.**

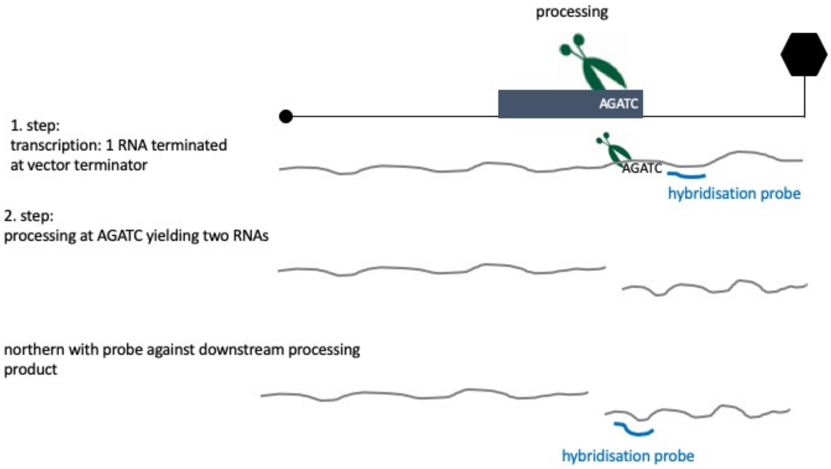

**B.**

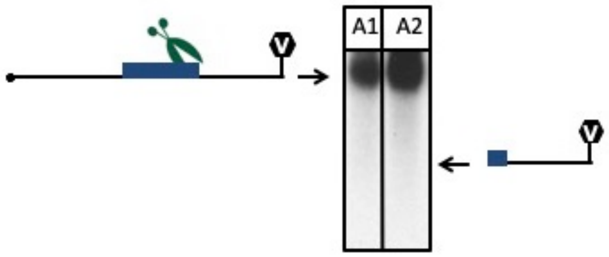

**C.**

**C. Legends to Supplementary Tables 1-4**

**Supplementary Table 1. Data for all transcription termination sites found with IE-PC.** This table contains information corresponding to TTS detected using IE-PC based on the +TEX data (identification of TTS). Columns are as follows:

**Block General information: #chr:** shows on which chromosome the TTS or RNA 3´ end is located; **TTS:** position of TTS/ RNA 3´ end in genome; **strand**: shows whether the TTS/ RNA 3´ end is on the minus or the plus strand; **usTTS:** distance to closest upstream TTS/ RNA 3´ end, if -1 then there is no other TTS/ RNA 3´ end in between the closest upstream annotated 3´ end and the current TTS/ RNA 3´ end.

**Block Upstream gene: usGene:** distance to closest upstream annotated gene, if -1 there is no upstream gene or information; **usGeneType:** type of upstream gene, e.g. CDS, RNA. If no upstream gene: NA; **usGeneID:** ID of upstream gene as given in the annotation.

**Block TTS/ RNA 3´ end in coding or intergenic region: inGene:** 1 if TTS/ RNA 3´ end is located inside annotated region, 0 if TTS/ RNA 3´ end is located in intergenic regions; **usFive:** distance to closest upstream annotated 5´ end, -1 if no information; **inGeneType:** type of gene the TTS/ RNA 3´ end is located in or NA if intergenic TTS/ RNA 3´ end; **inGeneID:** ID of gene the TTS/ RNA 3´ end is located in or NA if intergenic TTS/ RNA 3´ end.

**Block Sequence motif: seqMotif:** 1 if there is a sequence motif with the TTS/ RNA 3´ end, 0 if not; **motifNum:** number of motif as in paper, 0 if no motif, 1 or 2 for motifs.

**Block 3´ UTR: completeCov:** 1 if complete 3´ UTR is continuously covered with reads from RNAseq, 0 if not; **3´ UTR length:** distance between TTS and closest upstream 3´ end, -1 if no information; **avMeanCov:** mean of average coverage of 3´ UTR by RNAseq reads.

**Block Antisense complete gene: antisenseGene:** 1 if TTS/ RNA 3´ end is located in antisense gene, 0 if not; **antisenseStart:** start position of antisense gene, -1 if no antisense gene; **antisenseEnd:** end position of antisense gene, -1 if no antisense gene; **antisenseID:** ID of antisense gene as given in annotation.

**Supplementary Table 2. MEME motif search with data from IE-PC for +TEX reads. A. Motif in coding regions and B. in intergenic regions.** Sequence motifs for TTS detected using MEME are listed MEME (see also Figure 8 in main manuscript). For each TTS 15 nt upstream and 5 nt downstream were taken and used as an input sequence for MEME. The table columns are: genome part (CHR, pHV1, pHV3, pHV4), TTS position, start of input region, end of input region, strand, start of motif in input region, p-value.

**Supplementary Table 3. RNA 3´ ends discovered using DSM.** The table displays all RNA 3´ ends found with DSM. Columns: chromosome: type of chromosome; RNA 3´ end: position of the RNA 3´ end; corresponding average_insert_length: insert length of corresponding reads; strand: lists whether the RNA 3´ end is located on the plus or minus strand.

**Supplementary Table 4. 3´ UTRs covered by RNAseq data for RNA 3´ end data.** This table contains 10 columns which are also contained in Supplementary Table 1. Here, only 3´ UTRs are listed that are confirmed by continuous coverage of RNAseq reads.
Columns are:
**Block General information:** #chr: shows on which chromosome the TTS or RNA 3´ end is located; TTS: position of TTS/ RNA 3´ end in genome; strand: shows whether the TTS/ RNA 3´ end is on the minus or the plus strand; usTTS: distance to closest upstream TTS/ RNA 3´ end, if -1 then there is no other TTS/ RNA 3´ end in between the closest upstream annotated 3´ end and the current TTS/ RNA 3´ end.
**Block Upstream gene:** usGene: distance to closest upstream annotated gene, if -1 there is no upstream gene or information; usGeneType: type of upstream gene, e.g. CDS, RNA. If no upstream gene: NA; usGeneID: ID of upstream gene as given in the annotation.
**Block 3´ UTR:** 3´UTRcompleteCov: if 3´ UTR is continuously covered with reads from RNAseq: 1, if not: 0; 3´ UTR length: distance between TTS and closest upstream 3´ end, -1 if no information; avMeanCov: mean of average coverage of 3´ UTR by RNAseq reads.

### D. Supplementary Tables 5-8

### Supplementary Table 5. Primers, synthetic DNA, plasmids and strains used.

#### a. Strains

| strain | genotype | reference |
|---|---|---|
| *E.coli* DH5a | $F^-$, φ80d*lacZ*ΔM15, Δ(*lacZYA-argF*)U169, *deoR*, *recA*1, *endA*1, *hsdR*17($r_K^-$, $m_K^+$), *phoA*, *supE*44,$\lambda^-$, *thi*-1, *gyrA*96, *relA*1 | Stratagene |
| *H. volcanii* H119 | DS70(ΔpHV2), Δ*pyrE*2*,* Δ*trpA,* Δ*leuB* | Allers *et al*., 2004 |
| *H. volcanii* HV55 | DS70(ΔpHV2), Δ*pyrE*2*,* Δ*trpA,* Δ*leuB,* ΔHVO_A0326 | this work |

#### b. Plasmids

| plasmid | | reference |
|---|---|---|
| pTA231 | ColE1 ori, f1 ori, lacZ, $Amp^R$, *trpA*, pHV2 ori | Allers et al., 2004 |
| pTA617 | ColE1 ori, f1 ori, lacZ, $Amp^R$, *pyrE2*, flanking regions HVO_A0326 | Large et al., 2007 |

#### c. Synthetic DNA and inserts for termination test

| name | sequence |
|---|---|
| **synthetic DNA** | |
| psyn overexpression cassette | GCGGCCGCGAGCTCGGATCCGAGAATCGAAACGCTTATAAGTGCCCCCCGGCTAGAGAGCTGCATATGCTCGAGTCTAGATGAGCACCTCTGGACCATCGCATTTTTCGGCGTAAGCTTGGTACCGATATCGAATTC |
| **inserts for termination tests** | |
| bgaHa Cterminal fragment | CTCGTGACAGAGTTACTGGACGCCGCCGGCGTCGAGTACACTGAGCGATTCCCGGACGGCGTGCGCGTGATGGAGCGCGACGGCTATACGTGGGCGCTTAACTTCACGAGCGACCCGGTGACGTTGACCGTCCCCGATTCCACCGGGTTCCTGCTCGGTGAGTCCACCGTCGACGCGTTCGATACCGCGGTACTCGACGGATCCATCCGAGGTGTCGGACTCGCGTCCGAGTGAGTCGGCACGGGACCACTAGTTCTAGAGCGGCCGCCACCGCGGTGGAGCTCCAGCTTTTGTTCCCTTTAGTGAGGGTTAATTGCGCGCTTGGCGCTCGAGATATAA |
| Control | GGATCCGGCCGAGGCTGGAATCGAATACGTTCGAATGGGGGAGTTCGCGTGGCGACGAATCGAACCGGAGCGAGGGACGTTCGATTTCGCGTGGTTAGACGAGACCGTCGAACTCATCGGGAAGTCGGATCC |
| TTS-A1 | GGATCCGCACGGCCACCTCGACCACATCGGTGCCATCTCCAAGCTCGCCCACCGCTACGACGCCCCCGTCGTGGCGACGCCCTTCACCATCGAACTGGT |

| | |
|---|---|
| | GAAACAGCAGATCGAAGGCGAGAACAAGGATC C |
| TTS-A2 | GGATCCCGGCGAGGTCGAAGACGCCTTCGACGC CGGCGTCGTCGAGACGGCCCACGCCAAAGAGCA GGCCGTCGCCTCCGCCTCCGAAGCCGCGAACCT CGTCCTCAAGATCGACGACATCATCGCGGGATCC |
| TTS-S12 | GGATCCCTGAAGATGTGTCCAACAACCAGCAAAC AGACGATGAAATCCACGAGGACCAGCTCCTTAAC TTCCTCGTCAACTCTCTTGACGAGGAAGTTGCTC TCTCACTCGCTGAAAACGCTGAACTGGATCC |
| TTS-ftsZ2 | GGATCCGACGGCGGCCGCGACGAAGTCGAGAA GAACAACGGTCTCGACGTCATCCGGTAA CGCCCTGTCCGACCCGCGCGGACGCCGCCGCG CCCGTCGGGTTTCGTGGGTTCTTTTTCG CTGACGTGGATCC |

**d. Primers**

| **name** | **sequence** |
|---|---|
| ***bga*HDOprobe** | |
| bgaHKODO-for | GTCGGCACGGGACCACTC |
| bgaHKODO-rev | GACGGTGTACACGTGGAG |
| **bgaHa C-terminal fragment** | |
| bgaHatermi fw | CTCGTGACAGAGTTACTGGACGCC |
| bgaHatermi rev | TTATATCTCGAGCGCCAAGCGCGCAATTAAC |
| **terminator fragments** | |
| BamHI con fw | TTATATGGATCCGGCCGAGGCTGGAAT |
| TTS-A1fw | TTATAA GGATCCGCACGGCCACCTCGAC |
| TTS-A1rev | TTATAA GGATCC CTTGTTCTCGCCTTCG |
| TTS-A2fw | TTATAAGGATCCCGGCGAGGTCGAAGAC |
| TTS-A2rev | TTATAAGGATCCCGCGATGATGTCGTC |
| TermTestS12 fw | TTATATGGATCCCTGAAGATGTGTCCAAC |
| TermTestS12 rev | TTATATGGATCCAGTTCAGCGTTTTCAGC |
| TTS-ftsz fw | TTATATGGATCCGACGGCGGCCGCGACGAAGTC |
| TTS-ftsZ rev | TTATATGGATCCACGTCAGCGAAAAAGAACC |
| con2rev | ATATAAGGATCCGACTTCCCGATGAGTTC |
| **probes for northerns** | |
| TermiVectorrev | GTCGAGTACCGCGGTATCGAAC |
| BetaHinten1 | CGAGGTGTCGGACTCGCGTCC |
| BetaHinten2 | CGAGCGCCAAGCGCGCAATTAACC |

**Supplementary Table 6. Total reads, mapped and unmapped reads.** Three different types of RNAseq data were obtained: -TEX, +TEX and transcriptome data, each with three replicas. The number of reads, uniquely mapped reads and unmapped reads is listed as well as the percentage of mapped reads.

| sample | replica | total reads | uniquely mapped | un-mapped | % of mapped reads |
|---|---|---|---|---|---|
| -TEX | S1 | 42,814,712 | 40,848,267 | 1,966,445 | 95 |
| | S2 | 36,105,224 | 34,466,376 | 1,638,848 | 95 |
| | S3 | 47,288,670 | 45,649,822 | 1,638,848 | 97 |
| +TEX | S1 | 52,511,146 | 46,773,468 | 5,737,678 | 89 |
| | S2 | 32,123,046 | 29,191,284 | 2,931,762 | 91 |
| | S3 | 37,612,794 | 34,786,898 | 2,825,896 | 92 |
| transcriptome | S1 | 44,560,765 | 42,626,326 | 1,934,439 | 96 |
| | S2 | 42,013,751 | 40,447,008 | 1,566,743 | 96 |
| | S3 | 39,544,091 | 38,733,269 | 810,822 | 98 |

**Supplementary Table 7. Number of RNA 3´ ends determined using DSM.** The *Haloferax* genome consists of the main chromosome (2,848 kb) and three chromosomal plasmids pHV1, pHV3 and pHV4 (85, 438 and 636 kb, respectively)[22]. RNA 3´ ends are listed for each chromosome separately. In addition, the table shows whether the RNA 3´ ends are located in coding or in intergenic regions.

| chromosome | intergenic | coding | total |
|---|---|---|---|
| main | 2,046 | 378 | 2,424 |
| pHV1 | 70 | 22 | 92 |
| pHV3 | 204 | 18 | 222 |
| pHV4 | 353 | 64 | 417 |
| total | 2,673 | 482 | 3,155 |

**Supplementary Table 8. Determination of termination efficiencies.** Results of quantification of northern blot signals for the different terminator constructs are listed.Total RNA was isolated from strains transformed with the respective reporter gene plasmid. After size separation and transfer, membranes were hybridised with a probe against the β-galactosidase mRNA and signals were detected using phosphorimaging plates. For each lane, signals for termination at the vector terminator and the insert terminator were quantified and their sum set to 100 %; in columns “% vector length” and “% insert length” percentages for each transcript are given per lane. The row indicated by the sample name in bold gives the mean value of the three replicas (samples 1-3) assessed for each terminator motif and the column “SD” gives the standard deviation.

| Sample | % vector terminator | % insert terminator | SD |
|---|---|---|---|
| Con -1 | 99,7 | 0,3 | |
| Con -2 | 99,5 | 0,5 | |
| Con -3 | 98,8 | 1,2 | |
| **Control** | **99,3** | **0,7** | **0,5** |
| FtsZ -1 | 3,7 | 96,3 | |
| FtsZ -2 | 6,1 | 93,9 | |
| FtsZ -3 | 5,0 | 95,1 | |
| **FtsZ** | **4,9** | **95,1** | **1,2** |
| A1 -1 | 90,6 | 9,4 | |
| A1 -2 | 87,6 | 12,4 | |
| A1 -3 | 87,9 | 12,2 | |
| **A1** | **88,7** | **11,3** | **1,7** |
| A2 -1 | 73,0 | 27,0 | |
| A2 -2 | 72,2 | 27,8 | |
| A2 -3 | 73,9 | 26,2 | |
| **A2** | **73,0** | **27,0** | **0,8** |
| S12 -1 | 41,7 | 58,3 | |
| S12 -2 | 36,7 | 63,3 | |
| S12 -3 | 42,3 | 57,7 | |
| **S12** | **40,2** | **59,8** | **3,1** |
